# Genetic determinants of endophytism in the *Arabidopsis* root mycobiome

**DOI:** 10.1101/2021.04.28.441743

**Authors:** Fantin Mesny, Shingo Miyauchi, Thorsten Thiergart, Brigitte Pickel, Lea Atanasova, Magnus Karlsson, Bruno Hüttel, Kerrie W. Barry, Sajeet Haridas, Cindy Chen, Diane Bauer, William Andreopoulos, Jasmyn Pangilinan, Kurt LaButti, Robert Riley, Anna Lipzen, Alicia Clum, Elodie Drula, Bernard Henrissat, Annegret Kohler, Igor V. Grigoriev, Francis M. Martin, Stéphane Hacquard

**Author notes:** Corresponding authors: F. M. Martin and S. Hacquard.

## Abstract

Roots of *Arabidopsis thaliana* do not engage in symbiotic associations with mycorrhizal fungi but host taxonomically diverse fungal communities that influence health and disease states. We sequenced the genomes of 41 fungal isolates representative of the *A. thaliana* root mycobiota for comparative analysis with 79 other plant-associated fungi. We report that root mycobiota members evolved from ancestors with diverse lifestyles and retained large repertoires of plant cell wall-degrading enzymes (PCWDEs) and effector-like small secreted proteins. We identified a set of 84 gene families predicting best endophytism, including families encoding PCWDEs acting on xylan (GH10) and cellulose (AA9). These genes also belong to a core transcriptional response induced by phylogenetically-distant mycobiota members in *A. thaliana* roots. Recolonization experiments with individual fungi indicated that strains with detrimental effects in mono-association with the host not only colonize roots more aggressively than those with beneficial activities but also dominate in natural root samples. We identified and validated the pectin degrading enzyme family PL1_7 as a key component linking aggressiveness of endophytic colonization to plant health.

## Main

Roots of healthy plants are colonized by a rich diversity of microbes that can modulate plant physiology and development. Root colonization by arbuscular mycorrhizal and ectomycorrhizal fungi play fundamental roles in shaping plant evolution, distribution and fitness worldwide^1–6^. In contrast, the physiological relevance of root-associated fungi that do not establish symbiotic structures, but retain the ability to colonize roots of asymptomatic plants in nature remains unclear. These fungal endophytes can either represent stochastic encounters or engage in stable associations with plant roots^7–11^. Re-colonization experiments with individual fungal isolates and germ-free plants revealed various effects of mycobiota members on host performance, ranging along the parasitism-to-mutualism continuum^12–15^. Importantly, the outcome of the interaction on plant health can be modulated by host genetics, host nutritional status, and local environmental conditions^16–19^. Here, we assessed how evolutionary and functional properties of a diverse set of cultured fungi that colonize roots of the model plant *A. thaliana* can explain evolution towards endophytism and observed outcomes on plant performance. Using comparative genomics and transcriptomics, combined with re-colonization and functional validation experiments with germ-free plants, we showed that fungal colonization capabilities and effects on plant performance are negatively linked. Furthermore, we characterized a key genetic determinant of endophytism, the polysaccharide lyase family 1_7 that links fungal colonization aggressiveness to plant health. We propose that repertoires of PCWDEs of the *A. thaliana* root mycobiota shape fungal endosphere assemblages and modulate host fitness.

## Results

### Cultured isolates are representative of wild

A. thaliana root mycobiomes Fungi isolated from roots of healthy *A. thaliana* represent either stochastic encounters or robust endosphere colonizers. From a previously established fungal culture collection obtained from surface-sterilized root fragments of *A. thaliana* and relative *Brassicaceae* species^15^, we identified 41 isolates that could be distinguished based on their rDNA ITS1 sequences, representing 3, 27, and 37 different phyla, genera, and species of the fungal root microbiota, respectively (**Fig. 1a**). We first tested whether these phylogenetically diverse isolates were representative of naturally occurring root-colonizing fungi. Direct comparison with rDNA ITS1 sequence tags from a continental-scale survey of the *A. thaliana* root mycobiota^11^ revealed that most of the matching sequences were abundant (mean relative abundance, RA > 0.1 %, n = 30 / 41), prevalent (sample coverage > 50 %, n = 22 / 41), and enriched (root *vs.* soil, log2FC, Mann-Whitney-U test, *FDR* < 0.05, n = 26 / 41) in *A. thaliana* root endosphere samples at a continental scale (**Fig. 1a**). Quantitatively similar results were obtained using sequence data from the independent rDNA ITS2 locus (Spearman; Sample coverage: rs = 0.65, *P* < 0.01; RA: rs = 0.59, *P* < 0.01; **Fig. 1b**). The cumulative RA of the sequence tags corresponding to these 41 fungi accounted for 35 % of the total RA measured in root endosphere samples across European sites, despite the under-representation of abundant Agaricomycetes and Dothideomycetes taxa (**Fig. 1c**). We next assessed the worldwide distribution and prevalence of these fungal taxa across 3,582 root samples from diverse plants retrieved from the GlobalFungi database^20^. Continent-wide analysis revealed that the proportion of samples with positive hits was greater in Europe (sample coverage: up to 30 %, median = 4 %) than in North America (sample coverage: up to 10 %, median = 0.5 %), and largely insignificant in samples from other continents (**Fig. 1a**). Results indicate that most of the cultured *A. thaliana* root colonizing fungi are not stochastic encounters but reproducibly colonize plant roots across geographically distant sites irrespective of differences in soil conditions and climates.

**Fig. 1:**
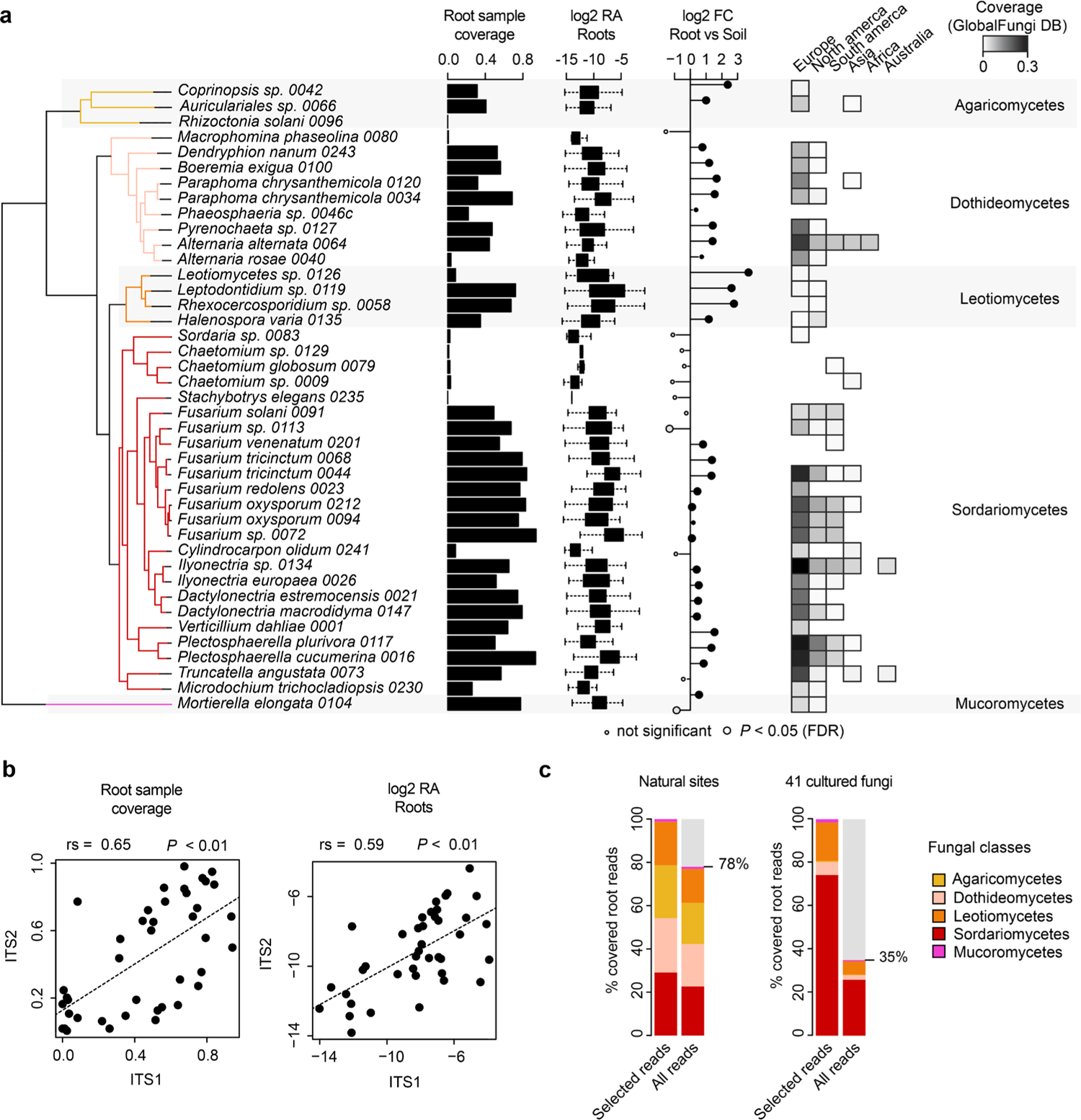
Prevalence and abundance profiles of 41 root-colonizing fungi across naturally occurring *A. thaliana* root mycobiomes. **a**, Species names and phylogenetic relationships among the 41 selected fungi. Estimated prevalence (i.e., root sample coverage, bar-plots), relative abundance (RA, log2 transformed, box-plots), and enrichment signatures (log2FC, circles) were calculated for each fungus based on data from a previously published continental-scale survey of the *A. thaliana* root mycobiota^11^. ITS1 tags from natural site samples were directly mapped against the reference ITS1 sequences of the selected fungi. Sample coverage in roots was computed based on n = 169 root samples and estimated RA were calculated for root samples having a positive hit only. Log2FC in RA between root (n = 169) and soil samples (n = 223) is shown based on the mean RA measured across samples and significant differences are indicated by circle sizes (Mann-Whitney-U test, FDR < 0.05). ITS1 sequence coverage measured across 3,582 root samples retrieved from the GlobalFungi^20^. Note that samples were analysed separately by continent. **b**, Correlation between root sample coverage (left panel) measured in ITS1 (n = 169) and ITS2 (n = 158) datasets for each of the 41 fungi (n = 41, Spearmańs rank correlation). Right panel: same correlation but based log2 RA values (n = 41, Spearmańs rank correlation). **c**, Distribution of root samples from the European transect across five classes of fungi, for ASVs (left panel) and the 41 fungal genomes (right panel). “Selected reads” refer to read distribution across the five fungal classes only, whereas “all reads” refer to read distribution across all sequenced reads.

### Root mycobiota members evolved from ancestors with diverse lifestyles

Given the broad taxonomic diversity of *A. thaliana* root mycobiota members, endosphere colonization capabilities may have evolved multiple times independently across distinct fungal lineages. We sequenced the above-mentioned 41 fungal genomes using PacBio long-read sequencing and annotated them with the support of transcriptome data (**Methods**), resulting in high-quality genome drafts (number of contigs: 9 - 919, median = 63; L50: 0.2 - 9.1 Mbp, median = 2.3 Mbp; **Supplementary Table 1**). Genome size varied between 33.3 and 121 Mb (median = 45 Mbp) and was significantly correlated with the number of predicted genes (number of genes: 10,414 - 25,647, median = 14,777, Spearman rs = 0.92, *P* = 3.82e-17) and the number of transposable elements (Spearman rs = 0.86, *P* = 4.13e-13) (**Extended Data Fig. 1**). A comparative genome analysis was conducted with 79 additional representative plant-associated fungi, selected based on their previously-described lifestyles^21^ (plant pathogens, soil/wood saprotrophs, ectomycorrhizal symbionts, ericoid mycorrhizal symbionts, orchid mycorrhizal symbionts and endophytes^17, 19, 22–27^) and close phylogenetic relationship to the strains we sequenced (**Fig. 2a**, **Extended Data Fig. 2, Supplementary Table 2**). To decipher potential evolutionary trajectories within this large fungal set, we first defined presence/absence of gene families in the 120 fungal genomes based on orthology prediction (N = 41,612; OrthoFinder^28^) and subsequently predicted the ancestral genome content using the Sankoff parsimony method (GLOOME^29^). Next, we trained a Random Forests classification model linking presence/absence of gene families to lifestyles, resulting in a lifestyle prediction accuracy of R^2^ = 0.56 (**Methods**). Although this classifier cannot confidently assign a single lifestyle to one genome content, it can be used to estimate lifestyle probabilities, and can reveal potential evolutionary trajectories when applied to Sankoff-predicted ancestral genomic compositions (see pie charts, **Fig. 2a**). This probabilistic approach corroborated that recent ancestors of the beneficial root endophyte *Colletotrichum tofieldiae* were likely pathogenic^17^, whereas those of beneficial Sebacinales - like those of ectomycorrhizal Agaricomycetes - were predicted to be saprotrophs^13, 30^. According to the classifier’s predictions, Agaricomycetes and Mortierellomycetes in *A. thaliana* mycobiota likely derive from soil saprotrophs, while the root-associated lifestyle seems ancestral in the classes Dothideomycetes, Leotiomycetes and Sordariomycetes, which emerged more recently^31^ (**Fig. 2a**). Therefore, *in planta* accommodation of *A. thaliana* root mycobiota members occurred multiple times, independently during evolution, as these fungi evolved from ancestors with diverse lifestyles.

**Fig. 2:**
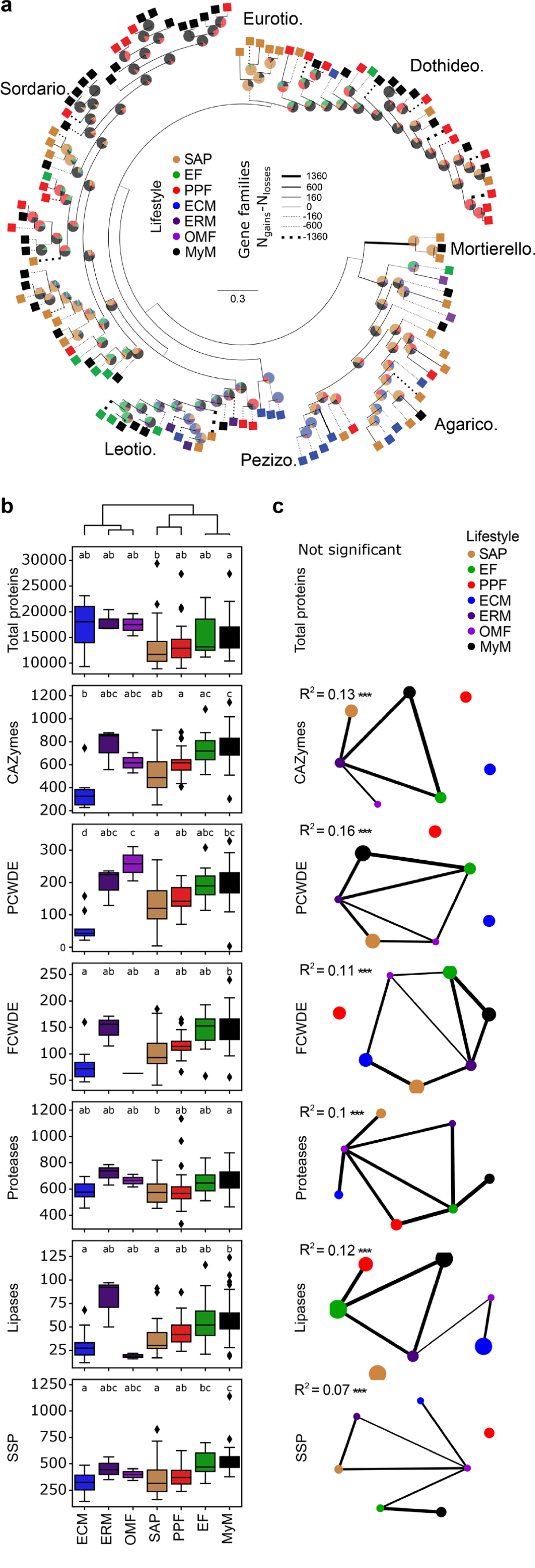
Ancestral relationships and trait convergence across root-colonizing fungal endophytes. **a**, Lifestyle-annotated whole genome phylogeny of the 41 selected mycobiota members (MyM, black) and 79 published fungal genomes (SAP: Saprotrophs, EF: Endophytic Fungi, PPF: Plant Pathogenic Fungi, ECM: Ectomycorrhiza, ERM: Ericoid Mycorrhiza, OMF: Orchid Mycorrhizal Fungi). Pie charts on ancestor nodes show lifestyle probabilities of each ancestor, as identified by a Random Forest model trained on genome compositions in gene families (R^2^ = 0.56). Branch width is proportional to the gene family gains-losses difference (N_gains_ – N_losses_). Line is dotted when this difference is negative. **b**, Genomic counts of genes involved in fungal-host/environment associations (CAZymes: Carbohydrate-Active Enzymes, PCWDEs: Plant Cell-Wall Degrading Enzyme, FCWDEs: Fungal Cell-Wall Degrading Enzyme, SSPs: Small Secreted Proteins; PCWDEs and FCWDEs are CAZyme subsets). Boxes are grouped according to UPGMA hierarchical clustering on mean counts over the different categories. ANOVA-statistical testing (Counts∼PhylogenyPCs+Lifestyle, **Methods**) identified both phylogeny and lifestyles as having an effect on genomic contents, letters result from post-hoc TukeyHSD testing. **c**, Networks showing the results of a PERMANOVA-based comparison of gene repertoires (JaccardDistances∼Phylogeny+Lifestyle). Networks for each category are labelled with Lifestyle R^2^ values. *** *P* < 0.001 (see **Extended Data Fig. 4**). Lifestyles are connected if their gene compositions are not significantly different. Node size is proportional to the area of one lifestyle’s ordination ellipse on a Jaccard-derived PCoA plot, and reflects the intra-lifestyle variability. Edge weights and widths are inversely proportional to the distance between ordination ellipse centroids.

### Functional overlap in genomes of root mycobiota members and endophytes

Isolation of mycobiota members from roots of healthy plants prompted us to test whether their gene repertoires more extensively resemble those of mycorrhizal symbionts, known endophytes, saprotrophs or pathogens. While the genomes of ectomycorrhizal fungi were shown to be enriched in transposable elements^32, 33^, the percentage of these elements remained low in genomes of root mycobiota members (0.69 % - 28.43 %, median = 5.44 %, **Extended Data Fig. 3**). We predicted genes known to play a role in fungal-host interactions (**Methods**), including those encoding carbohydrate-active enzymes (CAZymes), proteases, lipases and effector-like small secreted proteins (SSPs^34^), and then assessed differences in repertoire diversity across lifestyles (**Fig. 2b**). Unlike ectomycorrhizal fungi^32, 33^, but similarly to beneficial endophytes^16, 17, 19, 23, 27^, the genomes of root mycobiota members retained large repertoires of genes encoding plant cell wall-degrading enzymes (PCWDEs), SSPs and proteases (ANOVA-TukeyHSD, *P* < 0.05, **Fig. 2b**). Using Permutational multivariate analysis of variance (PERMANOVA based on Jaccard dissimilarity indices between genomes), we distinguished lifestyle from phylogenetic signals in gene repertoire composition (**Fig. 2c**, **Extended Data Fig. 4a**). This revealed that “lifestyle” significantly explained variation in gene repertoire composition (phylogeny: R^2^: 0.11 - 0.37, *P* < 0.05; lifestyle: R^2^: 0.07 - 0.16, *P* < 0.05, **Extended Data Fig. 4a**). Interestingly, the factor “lifestyle” explained the highest percentage of variance for PCWDE repertoires (phylogeny: R^2^ = 0.18; lifestyle: R^2^ = 0.16, **Extended Data Fig. 4a**), suggesting that these CAZymes likely play an important role in lifestyle differentiation. Further pairwise comparisons between lifestyle groups revealed that gene repertoire composition of root mycobiota members could not be differentiated from those of beneficial endophytes (*post-hoc* pairwise PERMANOVA, *P* > 0.05, **Fig. 2c**). Therefore, gene arsenals of *A. thaliana* root-colonizing fungi resemble those of endophytes more than saprotrophs, pathogens or mycorrhizal symbionts. Across the tested gene groups, the families which contribute the most in segregating genomes by lifestyles (**Extended Data Fig. 4b, Methods**) include two xylan esterases (CE1, CE5), two pectate lyases (PL3_2, PL1_4), one pectin methyltransferase (CE8) and one serine protease (S08A). Further analysis focusing on total predicted secretomes (**Extended Data Fig. 5, Extended Data Fig. 6a**) and CAZymes subfamilies (**Extended Data Fig. 6b**) confirmed strong genomic similarities between *A. thaliana* root mycobiota members and known endophytic fungi.

**Fig. 3:**
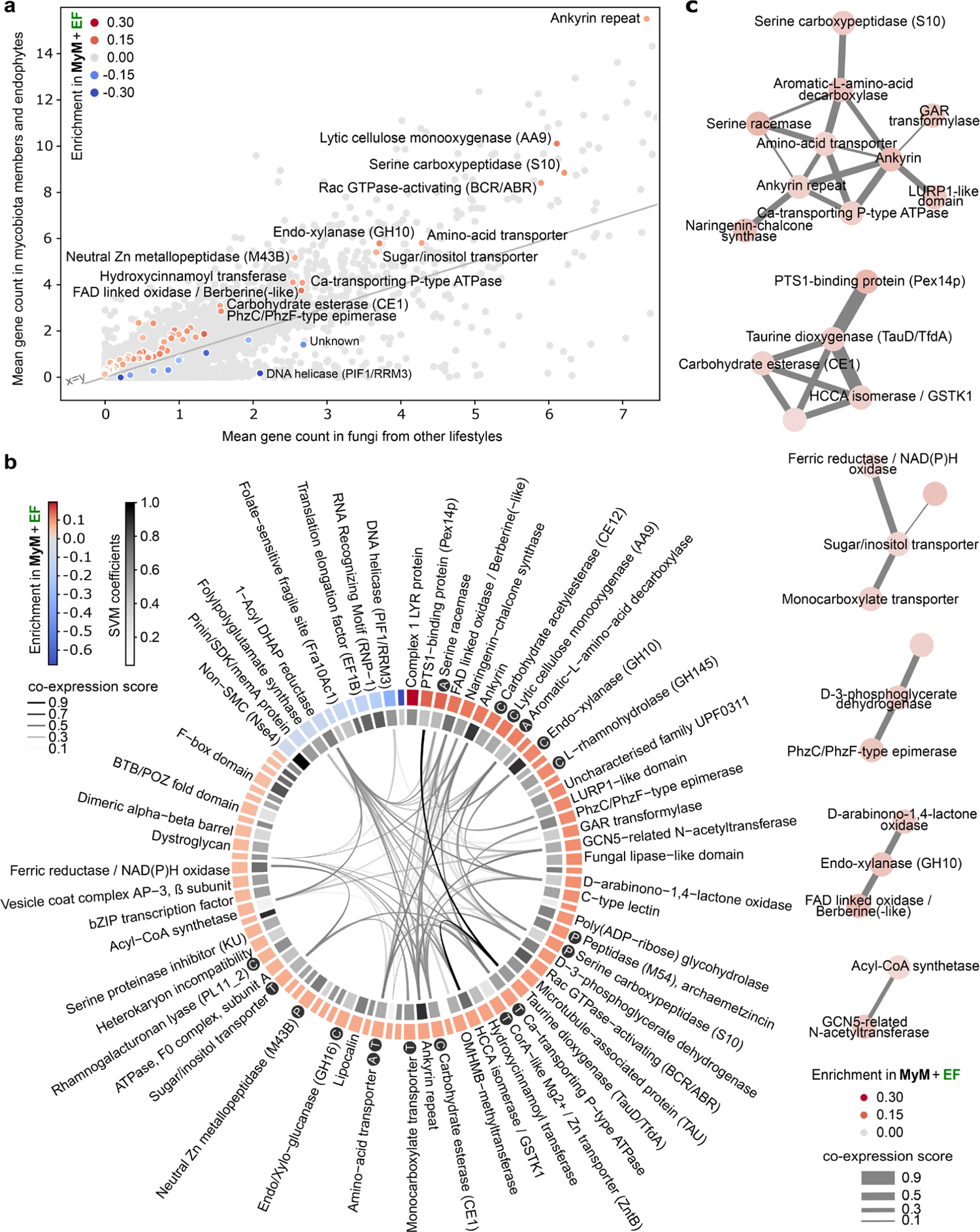
Minimal set of 84 gene families discriminating mycobiota members and endophytes from other lifestyles. **a**, Scatterplot showing the mean per-genome copy number of each orthogroup in mycobiota members and endophytes, in comparison to other lifestyles. Light grey: all 41,612 orthogroups. The 84 discriminant orthogroups identified by SVM-RFE (R^2^ = 0.8) are highlighted in a gradient of red or blue colours reflecting, respectively, enrichment or depletion in *A. thaliana* mycobiota members and endophytes (MyM + EF) compared to the other fungal lifestyles. **b**, Functional descriptions of the 84 discriminant orthogroups. This gene set is enriched in CAZymes (Fisher, *P* < 0.05 - labelled C) and also contains peptidases (labelled P), transporters (labelled T) and proteins involved in amino-acid metabolism (labelled A). The outer circle shows orthogroup enrichment/depletion as described in panel (a). The inner circle depicts the SVM coefficients, reflecting the contribution of each orthogroup to lifestyle differentiation. In the centre, links between orthogroups indicate coexpression of associated COG families in fungi (STRING database^37^). **c**, Coexpression network of gene families across published fungal transcriptomic datasets, built on discriminant orthogroups enriched in endophytes and mycobiota members and clustered with the MCL method.

**Fig. 4:**
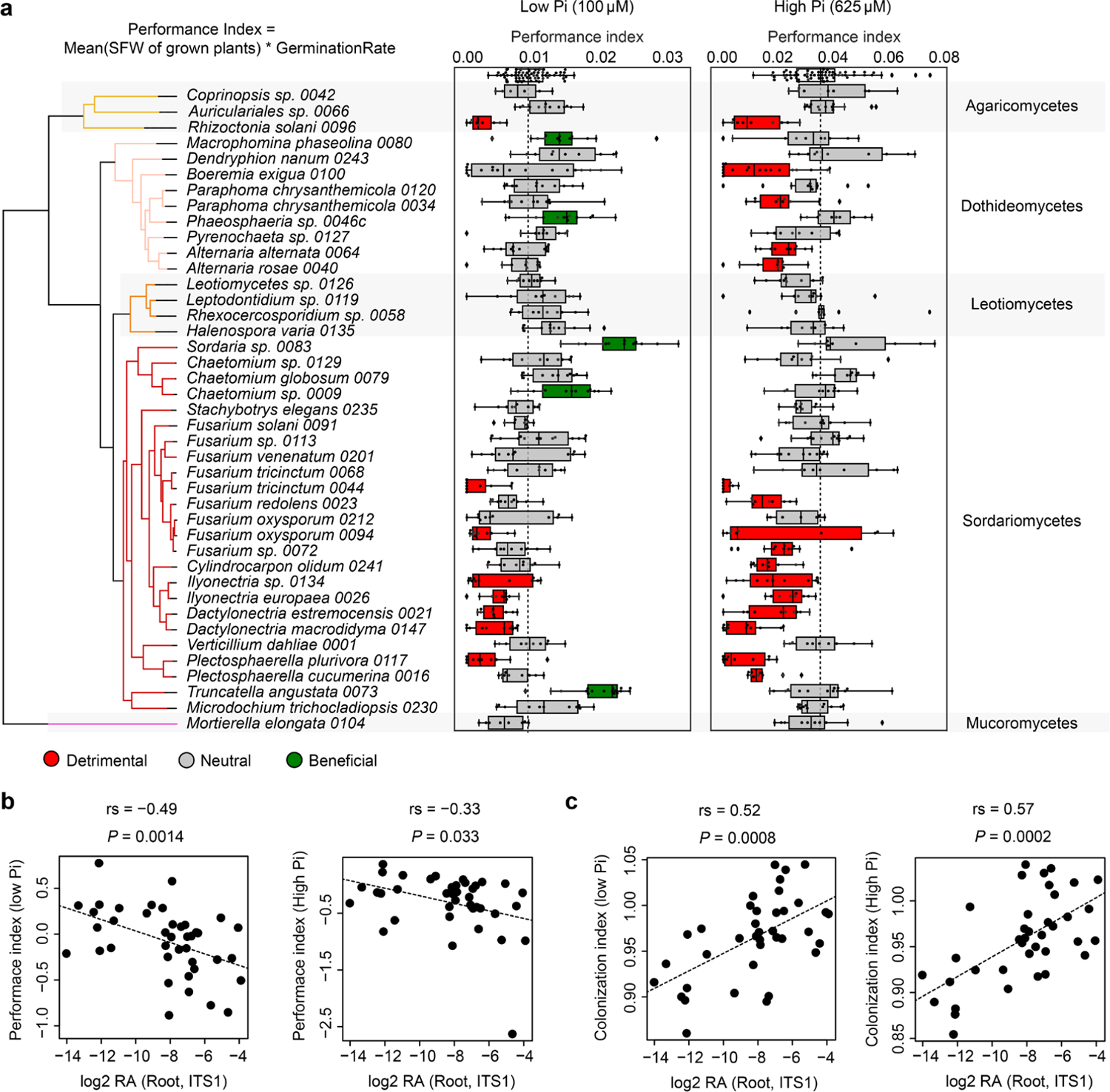
Linking fungal outcome on host performance with root colonization patterns. **a**, Performance indices (shoot fresh weights of 4-week-old plants normalized by germination rate) of *A. thaliana* plants recolonized with each of the 41 fungal strains on media containing low and high concentrations of orthophosphate (Pi). Differential fungal effects on plant performance were tested on both media with Kruskal-Wallis (*P* < 0.05) and beneficial and pathogenic strains were identified by a Dunn test against mock-treated plants (first row in boxplots). **b**, Spearman rank correlation of relative fungal abundances in root samples from natural populations^11^ (log2 RA, see Fig. 1a) with fungal effects on plant performance at low Pi (left) and high Pi (right) (Hedges standard effect sizes standardizing all phenotypes to the ones of mock-treated plants). **c**, Spearman rank correlation of relative fungal abundances in root samples from natural populations^11^ (log2 RA, see Fig. 1a) with fungal colonization indices measured by quantitative PCR in our plant recolonization experiments at low Pi (left) and high Pi (right).

**Fig. 5:**
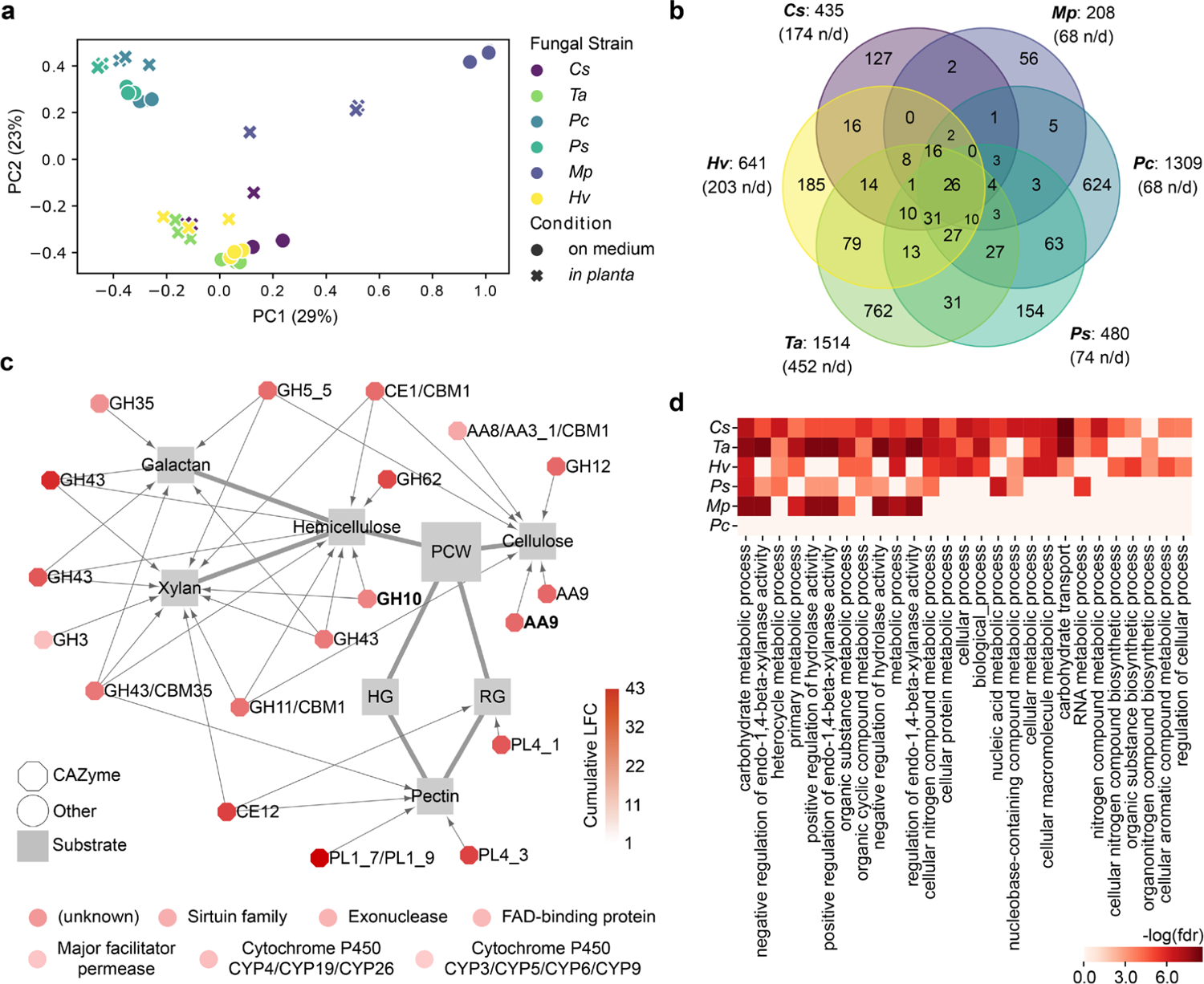
Comparative transcriptomics identified a core set of PCWDE-encoding genes induced in *A. thaliana* roots by diverse mycobiota members. **a**, PCoA plot of Bray-Curtis distances calculated on gene family read counts from fungal transcriptome data on medium and *in planta*. *Cs* = *Chaetomium sp. 0009*, *Mp* = *Macrophomina phaseolina 0080*, *Pc* = *Paraphoma chrysantemicola 0034*, *Ps* = *Phaeosphaeria sp. 0046c*, *Ta* = *Truncatella angustata 0073*, *Hv* = *Halenospora varia 0135.* b, Venn diagram showing the number of fungal gene families over-expressed *in planta*. Note the 26 families commonly over-expressed by all six fungi (n/d: non-displayed interactions). **c**, Commonly over-expressed gene families *in planta* (n = 26), which include 19 plant cell-wall degrading CAZymes (octagons) linked to their substrates, as described in literature^33, 64^. The two CAZyme families highlighted in bold were identified as potential determinants of endophytism (SVM-RFE, see Fig. 3a). The seven remaining (non-CAZyme) families are shown below the network. **d**, Individual GO enrichment analyses performed on the genes over-expressed *in planta vs.* on medium by each fungal strain (GOATOOLS, FDR < 0.05).

**Fig. 6:**
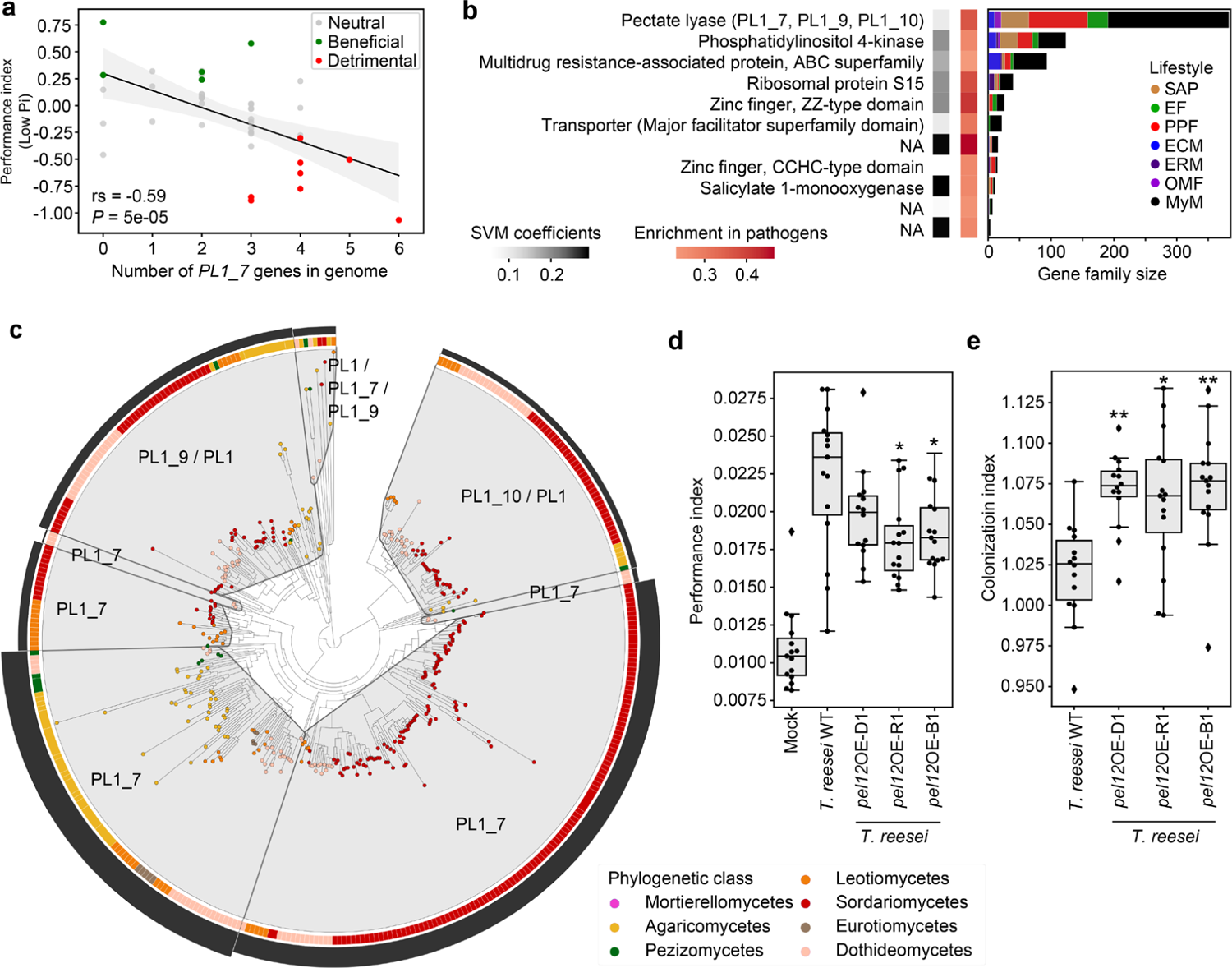
Genomic content in polysaccharide lyase PL1_7 links colonization aggressiveness to plant health. **a,** Spearman correlation between the number of genes encoding secreted PL1_7 in fungal genomes and the plant performance index at low Pi in recolonization experiments. **b**, Minimal set of 11 gene families discriminating detrimental from non-detrimental fungi at low Pi (SVM-RFE R^2^ = 0.88). The first vertical stripe shows the enrichment of these gene families in fungi identified as detrimental in recolonization experiments at low Pi, whereas the second shows the SVM coefficients, reflecting the contribution of each orthogroup to the separation of the two groups. Gene family sizes and representation in the different lifestyles are shown on the barplots in the context of the whole fungal dataset (n = 120). NA: no functional annotation. **c,** Protein family tree of the polysaccharide lyase orthogroup identified as essential for segregating detrimental from non-detrimental fungi in our SVM-RFE classification model. The tree was reconciled with fungal phylogeny and clustered into minimum instability groups by MIPhy^40^. Each group is labelled with its CAZyme annotation. The outer circle (black barplot) depicts the relative instabilities of these groups, suggesting two rapidly evolving PL1_7 groups in Sordariomycetes and Agaricomycetes. **d,** Plant performance indices resulting from plant recolonization experiments at low Pi, conducted with *Trichoderma reesei* QM9414 (WT) and three independent heterologous mutant lines (D1, R1, B1) overexpressing *pel12* from *Clonostachys rosea* (PL1_7 family^41^). Asterisks indicate significant difference to *T. reesei* WT, according to ANOVA and TukeyHSD test. **e,** Fungal colonization measured by qPCR in colonized roots at low Pi, conducted with *T. reesei* WT and three *pel12* overexpression mutant lines. Asterisks indicate significant difference to *T. reesei* WT, according to a Kruskal-Wallis and a Dunn test.

### Genomic traits of the endophytic lifestyle

To identify unique genetic determinants characterizing both known endophytes and *A. thaliana* root mycobiota members, the 120 genomes were mined for gene families whose copy numbers allow efficient segregation of these fungi (n = 50) from those with other lifestyles (n = 70). We trained a Support Vector Machines classifier with Recursive Feature Elimination (SVM-RFE) on the gene counts of orthogroups significantly enriched or depleted between these two groups (ANOVA, FDR < 0.05). A minimal set of 84 gene families that best segregated the two lifestyle groups was retained in the final SVM-RFE classifier (R^2^ = 0.80, **Fig. 3a** and **Supplementary Table 3**). These orthogroups can explain lifestyle differentiation independently from phylogenetic signal (PhyloGLM^35^ – 83 / 84, FDR < 0.05) and were significantly enriched in enzymes (i.e., GO *catalytic activity,* GOATOOLS^36^ FDR = 0.002, **Supplementary Table 3**) and in CAZymes (Fisher Exact Test, *P* = 0.03, data not shown). Notably, genes encoding PCWDEs acting on pectin (CE12, GH145, PL11), cellulose (AA9) and hemicellulose (i.e., xylan: GH10, GH16, CE1) were identified, together with others encoding peptidases, transporters and proteins involved in amino acid metabolism (**Fig. 3b** and **Supplementary Table 3**). These 84 gene families were analysed for co-expression in published fungal transcriptomic datasets (STRING^37^). An MCL-clustered co-expression network built on families enriched in known endophytes and *A. thaliana* mycobiota members revealed six clusters of co-expressed genes (**Fig. 3c**), including carbohydrate membrane transporters, and genes involved in carbohydrate metabolism (e.g., GH10) and amino acid metabolism. These functions are likely to be essential for root colonization and endophytism.

### Root colonization capabilities explain fungal outcome on plant growth

Root-colonizing fungi can span the endophytism-parasitism continuum but little is known regarding the genetic determinants that govern this delicate shift. We tested the extent to which the 41 fungi can modulate host physiology by performing binary interaction experiments with germ-free *A. thaliana* plants grown in two nutrient conditions under laboratory conditions (inorganic orthophosphate, Pi: 100 μM and 625 μM KH_2_PO_4_, **Fig. 4a**). We determined germination rate (GR) and shoot fresh weight (SFW) of five-week-old plants (n = 7,127) and used these parameters to calculate a plant performance index (PPI = SFW * GR, **Methods**). Under Pi-sufficient conditions, 39 % of the isolates (16 / 41) negatively affected host performance compared to germ-free control plants, whereas 61 % (25 / 41) had no significant effect on PPI (Kruskal-Wallis - Dunn Test, *adj. P* < 0.05, **Fig. 4a**). Fungal-induced change in PPI was significantly modulated by the nutritional status of the host, as depletion of bioavailable Pi in the medium was associated with a reduction in the number of fungi with pathogenic activities (20 %, 8 / 41) and an increase of those with beneficial activities (12 %, 5 / 41) (Kruskal-Wallis - Dunn Test, *adj. P* < 0.05, **Fig. 4a**). Notably, PPI measured for low and high Pi conditions was negatively correlated with strain RA in roots of European *A. thaliana* populations (Spearman, High Pi: rs = −0.33, *P* = 0.033; Low Pi: rs = −0.49, *P* = 0.0014, **Fig. 4b**), suggesting a potential link between the ability of a fungus to efficiently colonize roots and the observed negative effect on plant performance. Consistent with this hypothesis, fungal load measured by quantitative PCR in roots of five-week-old *A. thaliana* colonized by individual fungal isolates (**Extended Data Fig. 7a, b**), was positively correlated with fungal RA in roots of natural populations (Spearman, High Pi: rs = 0.57 *P* = 0.0002; Low Pi: rs = 0.52, *P* = 0.0008, **Fig. 4c**), and was also negatively linked with PPI outcome (Spearman, High Pi: rs = −0.44, *P* = 0.005, Low Pi: rs = −0.30, *P* = 0.057) (**Extended Data Fig. 7c, d**). Taken together, our results suggest that both nutritional constraints and controlled colonization abilities drive beneficial fungal-host associations.

A conserved set of CAZyme-encoding genes is induced *in planta* by diverse root mycobiota members.

We tested whether putative genomic determinants of endophytism defined above by a machine learning approach were part of a core response activated *in planta* by root mycobiota members. Six representative fungi from three different phylogenetic classes were selected for *in planta* transcriptomics on low Pi sugar-free medium: *Chaetomium sp.* 0009 (*Cs*), *Macrophomina phaseolina* 0080 (*Mp*), *Paraphoma chrysantemicola* 0034 (*Pc*), *Phaeosphaeria sp.* 0046c (*Ps*), *Truncatella angustata* 0073 (*Ta*), *Halenospora varia* 0135 (*Hv*). Confocal microscopy of roots grown in mono-association with these fungi highlighted similar colonization of root surfaces and local penetrations of hyphae in epidermal cells (**Extended Data Fig. 8**). After mapping of RNA-seq reads on genome assemblies (Hisat2^38^) and differential expression analysis (*in planta vs.* on medium, DESeq2^39^), significant log2 fold-change (log2FC) values were summed by orthogroups, allowing between-strain transcriptome comparisons (**Methods**). Transcriptome similarity did not fully reflect phylogenetic relationships since *Cs* and *Ta* (Sordariomycetes) clustered with *Hv* (Leotiomycete), whereas *Mp*, *Pc* and *Ps* (Dothideomycetes) showed substantial transcriptome differentiation (**Fig. 5a**). Although *in planta* transcriptional reprogramming was largely strain-specific, we identified a core set of 26 gene families that were consistently over-expressed by these unrelated fungi in *A. thaliana* roots (**Fig. 5b**). We observed a remarkable over-representation of genes coding for CAZymes acting on different plant cell wall components (i.e., 19 / 26, 73 %), including cellulose, xylan and pectin (**Fig. 5c**). This set was also significantly enriched in families previously identified as putative determinants of endophytism by our SVM-RFE classifier (Fisher exact test, *P* < 0.05), including AA9 and GH10 CAZyme families. Inspection of fungal genes over-expressed *in planta* by each strain (**Supplementary Table 4**), followed by independent GO enrichment analyses, corroborated that carbohydrate metabolic processes and xylanase activities were the most common fungal responses activated *in planta* (GOATOOLS, FDR < 0.05, **Fig. 5d**). Notably, we also observed important percentages of genes encoding effector-like SSPs induced *in planta* (9.8 % - 42.4 %, median = 21.6 %). Together, these enzymes and SSPs are likely to constitute an essential toolbox for *A. thaliana* root colonization and for fungal acquisition of carbon compounds from plant material. Analysis of corresponding *A. thaliana* root transcriptomes revealed that different responses were activated by the host as a result of its interaction with these six unrelated mycobiota members (**Extended Data Fig. 9, Supplementary Table 5**). Our data suggest that phylogenetically-distant mycobiota members colonize *A. thaliana* roots using a conserved set of PCWDEs and have markedly different impacts on their host.

### Polysaccharide lyase family PL1_7 as a key component linking colonization aggressiveness to plant health

We reported above a potential link between aggressiveness in root colonization and detrimental effect of fungi on PPI. To identify underlying genomic signatures explaining this link, we employed three different methods. First, inspection of diverse gene categories across genomes of beneficial, neutral, and detrimental fungi revealed significant enrichments in CAZymes (especially polysaccharide/pectate lyases, PLs) and proteases in the genomes of detrimental fungi (Low Pi conditions, Kruskal-Wallis and Dunn tests, **Extended Data Fig. 10a, b**). In these categories, three pectate lyases (PL1_4, PL1_7, PL3_2) and three peptidases (S08A, A01A, S10) contributed the most in segregating genomes by effect on plants (see the count in gene copy in **Extended Data Fig. 10c**). Second, multiple testing of association between secreted CAZyme counts (n = 199 families in total) and fungal effect on PPI identified the PL1_7 family as the only family significantly linked to detrimental effects (ANOVA, Bonferroni; Low Pi: *P* = 0.026; High Pi: not significant; **Fig. 6a**). Finally, an SVM-RFE classifier was trained on the gene counts of all orthogroups that were significantly enriched or depleted in genomes of detrimental *versus* non-detrimental fungi (ANOVA, FDR < 0.05). While this method failed at building a classifier to predict detrimental effects at high Pi (no families significantly enriched/depleted), it successfully predicted detrimental effects at low Pi with very high accuracy (R^2^ = 0.88). A minimal set of 11 orthogroups discriminating detrimental from non-detrimental fungi was identified (**Fig. 6b**, **Supplementary Table 6**), and includes gene families encoding membrane transporters, zinc-finger domain containing proteins, a salicylate monooxygenase and a PL1 orthogroup containing the aforementioned PL1_7 CAZyme subfamily and related PL1_9 and PL1_10 subfamilies. Further phylogenetic instability analysis based on duplication and mutation rates (MIPhy^40^) identified PL1_9 and PL1_10 as slow-evolving clades in the gene family tree (instability = 30.94 and 18.86 respectively, **Fig. 6c**), contrasting with most PL1_7 genes that were located in two rapidly-evolving clades (index = 85.30 and 66.12). Notably, genomic counts of PL1_7, but not PL1_9/10, remained significantly associated to detrimental host phenotypes after correction for the phylogenetic signal in our dataset (PhyloGLM^35^, FDR = 0.03). PL1_7 was also part of the core transcriptional response activated *in planta* by six non-detrimental fungi (**Fig. 5c**) and was enriched in mycobiota members and endophytes in comparison to saprotrophs and mycorrhizal fungi (**Extended Data Fig. 10d**). Therefore, degradation of pectin by root mycobiota members is likely crucial for penetration of — and accommodation in — pectin-rich *A. thaliana* cell walls. However, the remarkable expansion of this gene family in detrimental compared to non-detrimental fungi predicts a possible negative link between colonization aggressiveness and plant performance. To test this hypothesis, we took advantage of the *Trichoderma reesei* QM9414 strain (WT, PL1_7 free background) and its corresponding heterologous mutant lines over-expressing *pel12,* a gene from *Clonostachys rosea* encoding a PL1_7 pectate lyase with direct enzymatic involvement in utilization of pectin^41^. By performing plant recolonization experiments at low Pi with these lines, we observed that *T. reesei pel12*OE lines negatively affected PPI with respect to their parental strain (ANOVA and TukeyHSD test, *P* < 0.05 for two out of three independent overexpressing lines, **Fig. 6d**), and this phenotype was associated with a significant increase in fungal load in plant roots (Kruskal-Wallis and Dunn test, *P* < 0.05, **Fig. 6e**). Taken together, our data indicate that pectin-degrading enzymes belonging to the PL1_7 family are key fungal determinants linking colonization aggressiveness to plant health.

## Discussion

We report here that genomes of fungi isolated from roots of healthy *A. thaliana* harbour a remarkable diversity of genes encoding secreted proteins and CAZymes that have often been described for soil saprotrophs. However, given that these fungi were 1) isolated from surface-sterilized root fragments^15^, 2) enriched in plant roots *versus* surrounding soil samples at a continental scale^11^ (**Fig.1**), and 3) able to recolonize roots of germ-free plants (**Extended Data Figs 7** and **8**), it was not surprising that both the diversity and the composition of their gene repertoires resemble those of previously described endophytes^17, 19, 22^ (**Fig. 2**). Unlike the remarkable loss in PCWDE-encoding genes in the genomes of most ectomycorrhizal fungi^32, 33^, evolution towards endophytism in root mycobiota members is therefore not associated with genome reduction in saprotrophic traits, as previously suggested^16^. Using a machine learning approach, together with *in planta* transcriptomic experiments, we identified genes encoding CAZyme families AA9 (copper-dependent lytic polysaccharide monooxygenases, acting on cellulose chains) and GH10 (xylanase) as potential determinants of endophytism (**Figs 3** and **5**). Interestingly, these same families were strongly expanded in genomes of beneficial root mutualists belonging to *Serendipitaceae*^16, 42^ compared to mycorrhizal mutualists^32^ and might therefore represent key genetic components explaining adaptation to – and accommodation in – *A. thaliana* roots.

Although the 41 *A. thaliana* root mycobiota members were isolated from roots of healthy-looking plants, experiments in mono-associations with the host revealed a diversity of effects on plant performance, ranging from highly pathogenic to highly beneficial phenotypes (**Fig. 4**). These results are consistent with previous reports^12, 14, 15, 43^ and suggest that the pathogenic potential of detrimental fungal endophytes identified based on mono-association experiments with the host, is largely kept at bay in a community context by the combined action of microbiota-induced host defences and microbe-microbe competition at the soil-root interface^15, 44^. However, we observed that robust and abundant fungal colonizers of *A. thaliana* roots defined from a continental-scale survey of the root microbiota^11^ were dominated by detrimental fungi defined based on mono-association experiments with the host (**Fig. 4**). Based on quantitative PCR data, we also observed that fungi with beneficial activities on plant health were colonizing roots less aggressively than those with detrimental activities, suggesting a potential link between fungal colonization capabilities, abundance in natural plant populations and plant health. These results suggest that maintenance of fungal load in plant roots is critical for plant health, and that controlled fungal accommodation in plant tissues is key for the maintenance of homeostatic plant-fungal relationships. This conclusion is indirectly supported by the fact that an intact innate immune system is needed for beneficial activities of fungal root endophytes^16, 18^. Our results therefore suggest that the most beneficial root mycobiota members are not necessarily the most abundant in roots of natural plant populations. In contrast, understanding how potential pathogens can dominate the endospheric microbiome of healthy plants is key for predicting disease emergence in natural plant populations^45, 46^.

To identify genetic determinants explaining the link between colonization aggressiveness and detrimental effect on plant performance, we used different association methods that all converged into the identification of the CAZyme subfamily PL1_7 as one of the potential underlying determinants of this trait. Proteins from the PL1_7 family were previously characterized in different *Aspergillus* species as metabolizing pectate by eliminative cleavage of (1 -> 4)-α-D-galacturonan^47, 48^ (EC <u>4.2.2.2</u>).

Notably, primary cell walls of *A. thaliana* are enriched with pectin compared to those of monocotyledonous plants, which contain more hemicellulose and phenolics^49, 50^. Therefore, repertoire diversity in pectin-degradation capabilities is likely key for penetration and accommodation in pectin-rich *A. thaliana* cell walls. This is corroborated by the observation that non-detrimental fungal endophytes were also shown to consistently induce expression of this gene family *in planta* during colonization of *A. thaliana* roots (**Fig. 5**). However, re-inspection of previously published transcriptomic data indicated that genes encoding PL1_7 were induced more extensively *in planta* by the fungal root pathogen *Colletotrichum incanum* compared to that of its closely relative beneficial root endophyte *Colletotrichum toffeldiae*^17^. Therefore, differences in expression and diversification of this gene family are potential contributors to the differentiation between detrimental and non-detrimental fungi in the *A. thaliana* root mycobiome, especially since *Arabidopsis* cell-wall composition is a determinant factor for its disease resistance^51, 52^. Notably, expansion of the PL1_7 gene family was observed in plant pathogens but also in the biocontrol fungus *C. rosea* (Sordariomycetes, Hypocreales), a fungal species with mycoparasitic and plant endophytic capacity^53, 54^ that is phylogenetically closely related to multiple of our root mycobiota members. Genetic manipulation of the *C. rosea pel12* gene revealed a direct involvement of the protein in pectin degradation, but not in *C. rosea* biocontrol towards the phytopathogen *Botrytis cinerea*^41^. Here, we showed that heterologous overexpression of *C. rosea pel12* in *T. reesei* does not only increase its root colonization capabilities, but also modulates fungal impact on plant performance. We therefore conclude that a direct link exists between expression/diversification of PL1_7-encoding genes in fungal genomes, root colonization aggressiveness, and altered plant performance. Our results suggest that evolution of fungal CAZyme repertoires modulates root mycobiota assemblages and host health in nature.

## Methods

### Selection of 41 representative fungal strains

The 41 *A. thaliana* root mycobiota members were previously isolated from surface-sterilized root segments of *A. thaliana* and the closely related Brassicaceae species *Arabis alpina* and *Cardamine hirsuta,* as previously described^15^. Notably, this culture collection derived from fungi isolated from the roots of plants grown in the Cologne Agricultural Soil under greenhouse conditions, or from natural *A. thaliana* populations from two sites in Germany (Pulheim, Geyen) and on site in France (Saint-Dié, Vosges)^15^ (**Supplementary Table 1**).

### ITS sequence comparison with naturally occurring root mycobiome

Comparison of fungal ITS1 and ITS2 sequences with corresponding sequence tags from a European-scale survey of the *A. thaliana* mycobiota (17 European sites^11^) was carried out. For all 41 Fungi, sequences of the internal transcribed spacer 1 and 2 (ITS1 / ITS2) were retrieved from genomes (https://github.com/fantin-mesny/Extract-ITS-sequences-from-a-fungal-genome) or, in the cases where no sequences could be found, via Sanger sequencing (4 of 41). All ITS sequence variants were directly aligned to the demultiplexed and quality filtered reads from previously-published datasets^11^ using USEARCH at a 97% similarity cut-off^55^. A count table across all samples was constructed using the results from this mapping and an additional row representing all the reads that did not match any of the reference sequences was added. This additional row was based on the count data from the amplicon sequence variant (ASV) analysis from the original study, whereas the read counts from the new mapping were subtracted sample wise. To have coverage-independent information on the relative abundance (RA) of each fungus, we calculated RA only for the root samples where the respective fungi were found (RA > 0.01%). The sample coverage was calculated across all root samples (> 1,000 reads, n = 169). Enrichment in roots was calculated for all root and soil samples (> 1,000 reads, n = 169 / n = 223) using the Mann-Whitney-U test (FDR < 0.05). In order to estimate the presence of the 41 fungi across worldwide collected samples, we used the GlobalFungi database^20^ (https://globalfungi.com/, version August 2020). The most prevalent ITS1 sequences from each genome were used to conduct a BLAST search on the website. Sample metadata for the best matching representative species hypothesis sequences were then used to determine the global sample coverage. Appearance across samples from type ‘root’ was counted for each fungus and compared to the total number of root samples for each continent.

### Whole genome sequencing and annotation

Forty-one fungal isolates from a previously-assembled culture collection^15^ were revived from 30 % glycerol stocks stored at −80 °C. Genomic DNA extractions were carried out from mycelium samples grown on Potato extract Glucose Agar (PGA) medium, with a previously-described modified cetyltrimethylammonium bromide protocol^32^. Genomic DNA was sequenced using PacBio systems. Genomic DNA was sheared to 3 kb, > 10 kb, or 30 kb using using Covaris LE220 or g-Tubes or Megaruptor3 (Diagenode). The sheared DNA was treated with exonuclease to remove single-stranded ends and DNA damage repair mix followed by end repair and ligation of blunt adapters using SMRTbell Template Prep Kit 1.0 (Pacific Biosciences). The library was purified with AMPure PB beads and size selected with BluePippin (Sage Science) at > 10 kb cutoff size. Sequencing was done on PacBio RSII or SEQUEL machines. For RSII sequencing, PacBio Sequencing primer was annealed to the SMRTbell template library and sequencing polymerase was bound to them. The prepared SMRTbell template libraries were sequenced on a Pacific Biosciences RSII or Sequel sequencers using Version C4 or Version 2.1 chemistry and 1 x 240 or 1 x 600 sequencing movie run times, respectively. The genome assembly was generated using Falcon^56^ with mitochondria-filtered preads. The resulting assembly was improved with finisherSC, and polished with either Quiver or Arrow. Transcriptomes were sequenced using Illumina Truseq Stranded RNA protocols with polyA selection (http://support.illumina.com/sequencing/sequencing_kits/truseq_stranded_mrna_ht_sample_prep_kit.h tml) on HiSeq2500 using HiSeq TruSeq SBS sequencing kits v4 or NovaSeq6000 using NovaSeq XP v1 reagent kits, S4 flow cell, following a 2 x 150 indexed run recipe. After sequencing, the raw fastq file reads were filtered and trimmed for quality (Q6), artifacts, spike-in and PhiX reads and assembled into consensus sequences using Trinity^57^ v2.1.1. The genomes were annotated using the JGI Annotation pipeline^58^. Species assignment was conducted by extracting ITS1 and ITS2 sequences from genome assemblies, performing a similarity search against the UNITE database^59^ and a phylogenetic comparison to fungal genomes on MycoCosm^58^ (https://mycocosm.jgi.doe.gov).

### Comparative genomics dataset

In addition to our 41 fungal isolates from *A. thaliana* roots, we used 79 previously published fungal genomes in a comparative genomics analysis (**Supplementary Table 2**). While 77 genomes and annotations were downloaded from MycoCosm, the genome assemblies of fungal strains *Harpophora oryzae* R5-6-1^23^ and *Helotiales sp.* F229^19^ were downloaded from NCBI (GenBank assembly accessions GCA_000733355.1 and GCA_002554605.1 respectively) and annotated with FGENESH^60^. Lifestyles were associated to each single strain by referring to the original publications describing their isolation, and consulting the FunGuild^21^ database with the species and genus names associated to each strain. Orthology prediction was performed on this dataset of 120 genomes by running OrthoFinder v2.2.7^28^ with default parameters. From this prediction, we used the generated orthogroups data, the species tree and gene trees. OrthoFinder was also run on our 41 newly-sequenced fungi to obtain a second species tree, for this subset.

### Predicting ancestral lifestyles

To identify gene family gains and losses events and obtain reconstruction of ancestral genomes using the Sankoff approach, GLOOME^29^ *gainLoss.VR01.266* was run using the species tree and presence/absence of each orthogroup in the 120 genomes. To associate a lifestyle to each reconstructed ancestral genome, a random forest classifier was trained on the presence/absence of each orthogroup in the 120 genomes and their associated fungal lifestyles. This was performed using the RandomForestClassifier(random_state=0) function of the Python library *sklearn*^61^. The accuracy of the model was estimated by a leave-one-out cross-validation approach, computed using the function cross_val_score(cv=KFold(n_splits=120)) in *sklearn*. Finally, the probabilities of ancestors to belong in each lifestyle category were retrieved using function predict_proba().

### Genomic feature analyses

Statistics of genome assemblies (i.e., N50, number of genes and scaffolds and genome size) were obtained from JGI MycoCosm, and assembly-stats (https://github.com/sanger-pathogens/assembly-stats). Genome completeness with single copy orthologues was calculated using BUSCO v3.0.2 with default parameters^62^. The coverage of transposable elements in genomes was calculated and visualized using a custom pipeline Transposon Identification Nominative Genome Overview (TINGO^63^). The secretome was predicted as described previously^34^. We calculated, visualized, and compared the count and ratio of total (present in the genomes) and predicted secreted CAZymes^64^, proteases^65^, lipases^66^, and small secreted proteins^34^ (SSPs) (< 300 amino acid) as a subcategory. We calculated the total count of the followings using total and predicted secreted plant cell-wall degrading enzymes (PCWDEs) and fungal cell-wall degrading enzymes (FCWDEs). Output files generated above were combined and visualised with a custom pipeline, Proteomic Information Navigated Genomic Outlook (PRINGO^33^). To compare the genomic compositions of the different lifestyle categories while taking into account phylogenetic signal, we first generated a matrix of pairwise phylogenetic distances between genomes using the function tree.distance() from package *biopython Phylo*^67^, then computed a principal component analysis using the PCA(n_components=2) function of *sklearn*^61^. Components PC1 and PC2 were then used to compare the per-genome numbers of CAZymes, proteases, lipases, SSPs, PCWDEs and FCWDEs in the different lifestyles with an ANOVA test and a TukeyHSD post-hoc test. R function *aov* was used with the following formula specifying the model: *GeneCount∼PC1+PC2+Lifestyle+PC1:Lifestyle+PC2:Lifestyle*.

Differences in subfamily composition of the groups of genes of interest were then carried out using a PERMANOVA-based approach (https://github.com/fantin-mesny/Effect-Of-Biological-Categories-On-Genomes-Composition). This approach relies on function adonis2() from R package *Vegan* (https://github.com/jarioksa/vegan) and post-hoc testing with function pairwise.perm.manova() from package *RVAideMemoire* (https://cran.r-project.org/web/packages/RVAideMemoire). We determined genes discriminating groups based on the principal coordinates of a regularized discriminant analysis calculated from the count of genes coding for CAZymes, proteases, lipases, and small secreted proteins, with R function rda(). We then used Vegan function scores() on the three first principal coordinates, and kept for each coordinate the top five high-loading gene discriminating groups.

### Determinants of endophytism

To identify a small set of orthogroups that best segregate endophytes and mycobiota members from fungi with other lifestyles, we standardized the orthogroup gene counts with function StandardScaler() from *sklearn*^61^. Then, orthogroups that are enriched or depleted in the fungi of interest were selected with function SelectFdr(f_classif, alpha=0.05) from *sklearn*. On this subset of orthogroups, we trained a Support Vector Machine classifier with Recursive Feature Elimination (SVM-RFE). This was performed with functions from *sklearn* SVC(kernel=’linear’) and RFECV(step=10, cv=KFold(n_splits=120, min_features_to_select=10), which implement a leave-one-out cross-validation allowing the estimation of the classifier accuracy at each step of the recursive orthogroup elimination. PhyloGLM models^35^ were built on the two groups of interest and orthogroup gene counts, with parameters btol=45 and log.alpha.bound=7, and the *logistic_MPLE* method. Further analysis of the gene families segregating fungi of interest from others (n = 84) was carried out by identifying a representative sequence of each orthogroup in our SVM-RFE model, and studying both its annotation and coexpression data in databases. To identify representative sequences, all protein sequences composing an orthogroup were aligned with FAMSA^68^ v1.6.1. Using HMMER^69^ v3.2.1, we then built a Hidden Markov Model (HMM) from this alignment with function hmmbuild, then ran function hmmsearch looking for the best hit matching this HMM within the proteins composing our orthogroup. We then considered this best hit as a representative sequence of the orthogroup and analyzed its annotation. GO enrichment analysis was performed by running GOATOOLS^36^ using the GO annotations associated to the representative sequences. To obtain coexpression data linking the orthogroups retained in our SVM-RFE model, we searched the String-db^37^ website for COG protein families matching our set of representative protein sequences in fungi. Each protein was associated to one COG (**Supplementary Table 3**), and coexpression data were downloaded. A coexpression network was then built on the families enriched in endophytes and mycobiota members (n = 73) and clustered with algorithm MCL (granularity = 5) using Cytoscape^70^ v3.7.2 and clusterMaker2^71^ v1.3.1.

### Plant recolonization experiments assessing the effect of each fungal strain on plant growth

*A. thaliana* seeds were sterilized 15 min in 70 % ethanol, then 5 min in 8 % sodium hypochlorite. After 6 washes in sterile double-distilled water and one wash in 10 mM MgCl_2_, they were stratified 5 to 7 days at 4 °C in the dark. Seed inoculation with fungal strains was carried out by crushing 50 mg of mycelium grown for 10 days on Potato extract Glucose Agar medium (PGA) in 1 ml of 10 mM MgCl_2_ with two metal beads in a tissue lyser, then adding 10 µM of this inoculum in 250 µl of seed solution for 5 min. Seeds were then washed twice with MgCl_2_ before seven were deposited on each medium-filled square Petri plate. Mock-inoculated seeds were also prepared by simple washes in MgCl_2_. The two media used in this study — 625 and 100 µM Pi — were previously-described^72^. They were prepared by mixing 750 µM MgSO_4_, 625 µM / 100 µM KH_2_PO_4_, 10,300 µM NH_4_NO_3_, 94,00 µM KNO_3_, 1,500 µM CaCl_2_, 0.055 µM CoCl_2_, 0.053 µM CuCl_2_, 50 µM H_3_BO_3_, 2.5 µM KI, 50 µM MnCl_2_, 0.52 µM Na_2_MoO_4_, 15 µM ZnCl_2_, 75 µM Na-Fe-EDTA, and 1,000 µM MES pH5.5, 0µM / 525 µM KCl, then adding Difco^TM^ Agar (ref. 214530, 1% final concentration), and finally adapting the pH to 5.5 prior to autoclaving. Plants were grown for 28 days at 21 °C, for 10 hours with light (intensity 4) at 19 °C and 14 hours in the dark in growth chambers. While roots were harvested and flash-frozen, shoot fresh weight was measured for each plant. To distinguish seeds that did not germinate from plants that could not develop because of a fungal effect, we introduced a per-plate plant performance index corresponding to the average shoot fresh weight of grown plants multiplied by the proportion of grown plants. In further correlation analyses, we used plant-performance indexes normalized to mock controls (standard effect sizes) using the Hedges’ *g* method^73^.

### Fungal colonization of roots assay

Frozen root samples (one per plate) were crushed and total DNA was extracted from them using a QIAGEN Plant DNEasy Kit. Fungal colonization of these root samples was then measured by quantitative PCR. For each sample, two reactions were conducted with primers ITS1F (5’-CTTGGTCATTTAGAGGAAGTAA-3’) and ITS2 (5’-GCTGCGTTCTTCATCGATGC-3’) which target the fungal ITS1 sequence, and two with primers UBQ10F (5’-TGTTTCCGTTCCTGTTATCT-3’) and UBQ10R (5’-ATGTTCAAGCCATCCTTAGA-3’) that target the *Ubiquitin10 A. thaliana* gene. Each reaction was performed by mixing 5 µl of iQ SYBR Green Supermix with 2 µl of 10 µM forward primer, 2 µl of 10 µM reverse primer and 1µl of water containing 1 ng template DNA. A BioRad CFX Connect Real-Time system was used with the following programme: 3 min of denaturation at 95 °C, followed by 39 cycles of 15 sec at 95 °C, 30 sec at 60 °C and 30 sec at 72 °C. We then calculated a single colonization index for each sample using the following formula: 2^(-averageCq(ITS1)/averageCq(UBQ10)).

### Confocal microscopy of root colonization by fungi

Roots of plants grown for 28 days in mono-association with fungi were harvested and conserved in 70 % ethanol. They were then rinsed in ddH2O, and stained with propidium iodide (PI) and wheat germ agglutinin conjugated to fluorophore Biotium CF®488 (WGA-CF488). This was carried out by dipping the root samples for 15 min in a solution of 20 µg / ml PI and 10 µg / ml WGA-CF488 buffered at pH 7.4 in phosphate-buffered saline (PBS). Samples were then washed in PBS and imaged with a Zeiss LSM700 microscope.

### Plant-fungi interaction transcriptomics

Dual RNAseq of six different plant-fungi interactions was carried out by performing plant recolonization experiments on our low Pi medium, as described above. Total roots per plates were harvested after 28 days in culture, flash frozen, and crushed in a tissue lyser, and then total RNA was extracted with a QIAGEN RNeasy Plant Mini kit. As a control condition, sterile Nucleopore Track-Etched polyester membranes were deposited on low Pi medium, then 10 µl drops of fungal inoculum (50 mg / ml of mycelium in 10 mM MgCl_2_) were placed on each one. The membranes were collected and processed as the root samples of our test condition. PolyA-enrichment was carried out on the RNA extracts, then an RNAseq library was prepared with the NEBNext Ultra™ II Directional RNA Library Prep Kit for Illumina (New England Biolabs). Sequencing was then performed in single read mode on a HiSeq 3000 system. RNAseq reads were trimmed using Trimmomatic^74^ and parameters TRAILING:20 AVGQUAL:20 HEADCROP:10 MINLEN:100. We then used HiSat2^38^ v2.2.0 to map the trimmed reads onto reference genomes. Six independent HiSat2 indexes were prepared, each based on the TAIR10 *Arabidopsis thaliana* genome and one of the six fungal genome assemblies of interest. We then performed six mappings, and counted the mapped reads using featureCounts^75^. RPKM (Reads Per Kilobase Million) values were computed from the featureCounts output. Differential gene expression analyses were then carried out on these counts using DESeq2^39^. log2FC values were corrected by shrinkage with the algorithm *apeglm*^76^. To compare the transcriptomes of the six different fungi, significant log2FC values were summed per orthogroup. For each orthogroup, we used annotation of the most representative sequence, as previously described. GO enrichment analyses were carried out with GOATOOLS^36^, using the MycoCosm^58^ GO annotation for fungi, and the TAIR annotation for *Arabidopsis thaliana*.

### Determinants of detrimental effects on plants and analysis of pectate lyases

Determinants of detrimental effects at low Pi were identified with the same method as previously described for determinants of endophytism/mycobiota: standard scaling of the orthogroup gene counts, then training of an SVM classifier with RFE and leave-one-out cross validation. Instability analysis was carried out by submitting the species tree generated by OrthoFinder^28^ to MIPhy^40^, together with the gene tree of our orthogroup of interest, with default parameters. PhyloGLM^35^ models were built on the two groups detrimental/non-detrimental and CAZyme gene counts, using our 41-genome species tree with default parameters and the *logistic_MPLE* method. *T. reesei* strain QM9414 and three heterologous overexpression lines of *pel12* generated as described previously^41^, were revived on PGA medium and then inoculated into seeds for plant recolonization experiments on low Pi medium as previously described.

## Statistics

Except for statistical methods described in the previous paragraphs, statistical testing was performed in *R v3.5.1*. Function aov() was used for ANOVA tests, and TukeyHSD post-hoc testing was performed using function TukeyHSD(). The non-parametric Kruskal-Wallis test was used by running function kruskal.test(), and the Dunn post-hoc test was performed with function DunnTest() from package DescTools (https://github.com/AndriSignorell/DescTools/).

## Reporting Summary

Further information on research design is available in the Nature Research Reporting Summary linked to this article.

## Data availability

Raw and processed genome sequencing data are available at MycoCosm (https://mycocosm.jgi.doe.gov/mycocosm/home) and GenBank accessions will be provided upon publication. Raw and processed transcriptomic data used in our differential gene expression analysis are available at Gene Expression Omnibus: GSE169629. Scripts used for data processing and analysis are available at https://github.com/fantin-mesny/Scripts-from-Mesny-et-al.-2021

## Acknowledgements

The sequencing project was funded by the U.S. Department of Energy (DOE) Joint Genome Institute, a DOE Office of Science User Facility, and supported by the Office of Science of the U.S. DOE under Contract No. DE-AC02-05CH11231 within the framework of CSP 1974 “1KFG: Deep-sequencing of ecologically-relevant Dikaria”. This work was supported by funds to S.Hac from a European Research Council starting grant (MICRORULES 758003), the ‘Priority Programme: Deconstruction and Reconstruction of the Plant Microbiota (SPP DECRyPT 2125)’ and the Cluster of Excellence on Plant Sciences (CEPLAS), both funded by the Deutsche Forschungsgemeinschaft. F.M. salary was covered by the DECRyPT 2125 programme. This research was also supported by the Laboratory of Excellence ARBRE (ANR-11-LABX-0002-01), the Region Lorraine, the European Regional Development Fund, and the Plant–Microbe Interfaces Scientific Focus Area in the Genomic Science Program, the Office of Biological and Environmental Research in the US DOE Office of Science (to F.M.M). M.K acknowledge funding from the SLU Centre for Biological Control (CBC). The Austrian Science Fund FWF project P30460-B32 is acknowledged for funding L.A.. We would like to thank Nathan Vannier for regular discussions and ideas about data analysis and method development. Finally, we thank Paul Schulze-Lefert, Ruben Garrido-Oter, Ryohei Thomas Nakano, Gregor Langen and Rozina Kardakaris for providing helpful comments regarding the manuscript or during departmental seminars and thesis advisory committee meetings.

## Contributions

S.Hac. and F.M.M. initiated, coordinated and supervised the project. Genomic analyses were performed by F.M., with help of S.M. who provided scripts and pipelines to annotate genomic features and generate figures describing the structure and composition of fungal genomes. Transcriptomic analyses were performed by F.M. T.T. re-analysed ITS amplicon sequencing data from natural site samples. F.M. performed all the experiments, with technical assistance from B.P.. L.A. and M.K. provided fungal mutant strains used for functional validation of PL1_7. K.W.B., S.Har., C.C., D.B., A.L., W.A., J.P., K.L., R.R., A.C., and I.V.G. sequenced, assembled and annotated the fungal genomes. E.D. and B.He. annotated CAZymes. B.Hü. prepared RNAseq libraries and sequenced the transcriptomes used for differential gene expression analysis. F.M. and S.Hac. wrote the manuscript, with input from S.M., A.K., and F.M.M.

## Extended data figures

**Extended Data Fig. 1:**
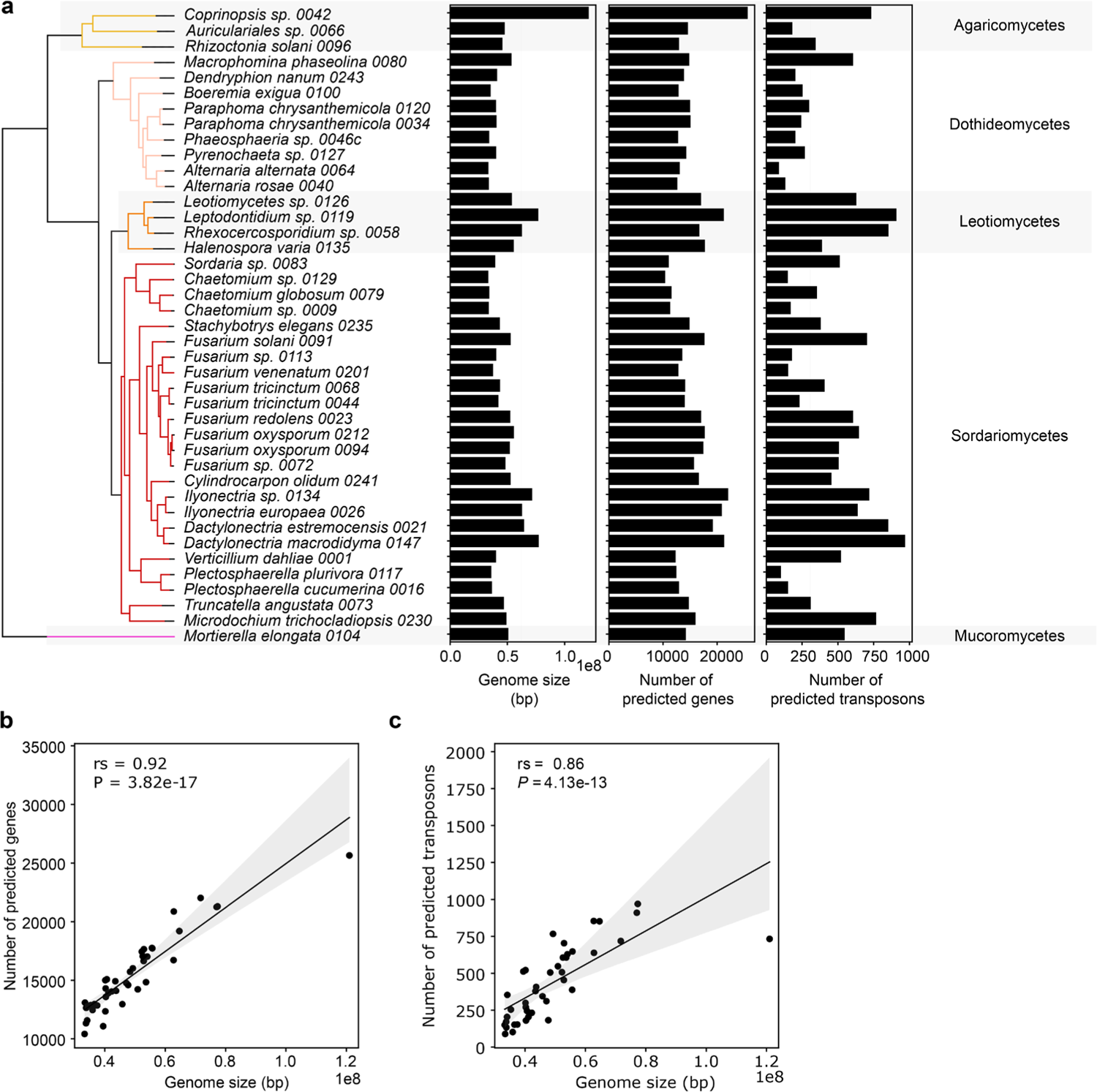
Link between genome size, number of genes and number of transposons across the 41 newly-sequenced fungal strains. **a**, Genome assembly size, number of predicted genes and number of identified transposons in the genomes of the 41 *A. thaliana* mycobiota members. **b**, Spearman’s correlation between genome size and number of predicted genes. **c**, Spearman’s correlation between genome size and number of predicted transposable elements.

**Extended Data Fig. 2:**
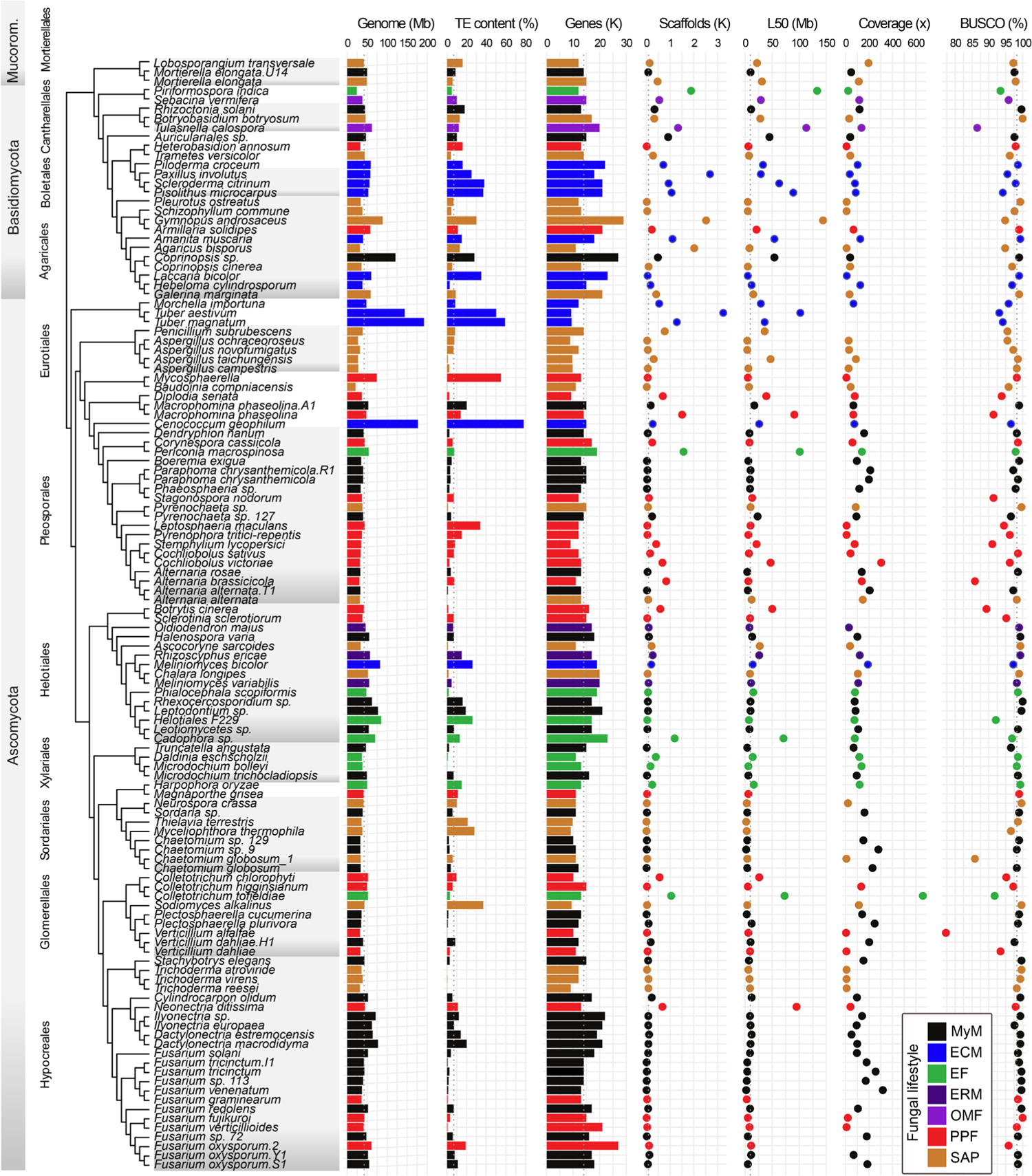
Genome sizes and properties of the 41 root mycobiota members, along with 79 previously published genomes used for comparative genomics. The species are in the evolutionary order. Fungal lifestyle is depicted in colour. Median values are depicted in dotted line. Genome: Genome size. TE content: The coverage of transposable elements in the genomes. Genes: The number of genes. Secreted: The number of theoretically secreted proteins (Methods). Scaffolds: The number of scaffolds. L50: N50 length. Coverage: Sequencing depth in fold. BUSCO: Genome completeness. SAP: Saprotrophs, EF: Endophytic Fungi, PPF: Plant Pathogenic Fungi, ECM: Ectomycorrhiza, ERM: Ericoid Mycorrhiza, OMF: Orchid Mycorrhizal Fungi, MyM *A. thaliana* mycobiota members.

**Extended Data Fig. 3:**
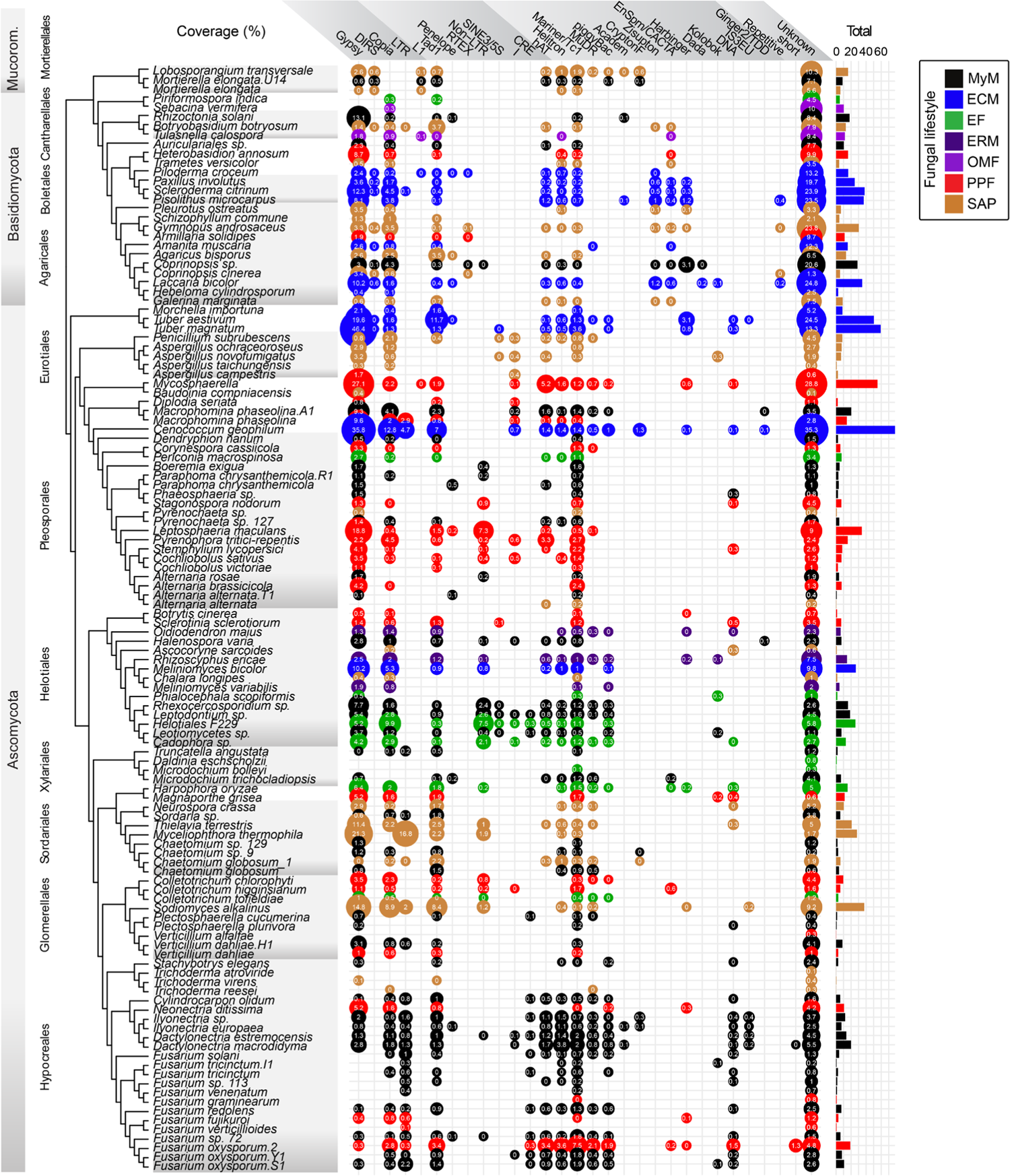
Genomic compositions in transposable elements across the fungal dataset. Coverage of transposable elements in 120 fungi analysed. LTR: Long terminal repeat retrotransposons. Non-LTR: Non-long terminal repeat retrotransposons. DNA: DNA transposons. Repetitive/short: Simple repeats. Unknown: Unclassified repeated sequences. The bubble size is proportional to the coverage of each of the transposable elements (shown inside the bubbles). The right bars show the total coverage per genome. SAP: Saprotrophs, EF: Endophytic Fungi, PPF: Plant Pathogenic Fungi, ECM: Ectomycorrhiza, ERM: Ericoid Mycorrhiza, OMF: Orchid Mycorrhizal Fungi, MyM: *A. thaliana* mycobiota members.

**Extended Data Fig. 4:**
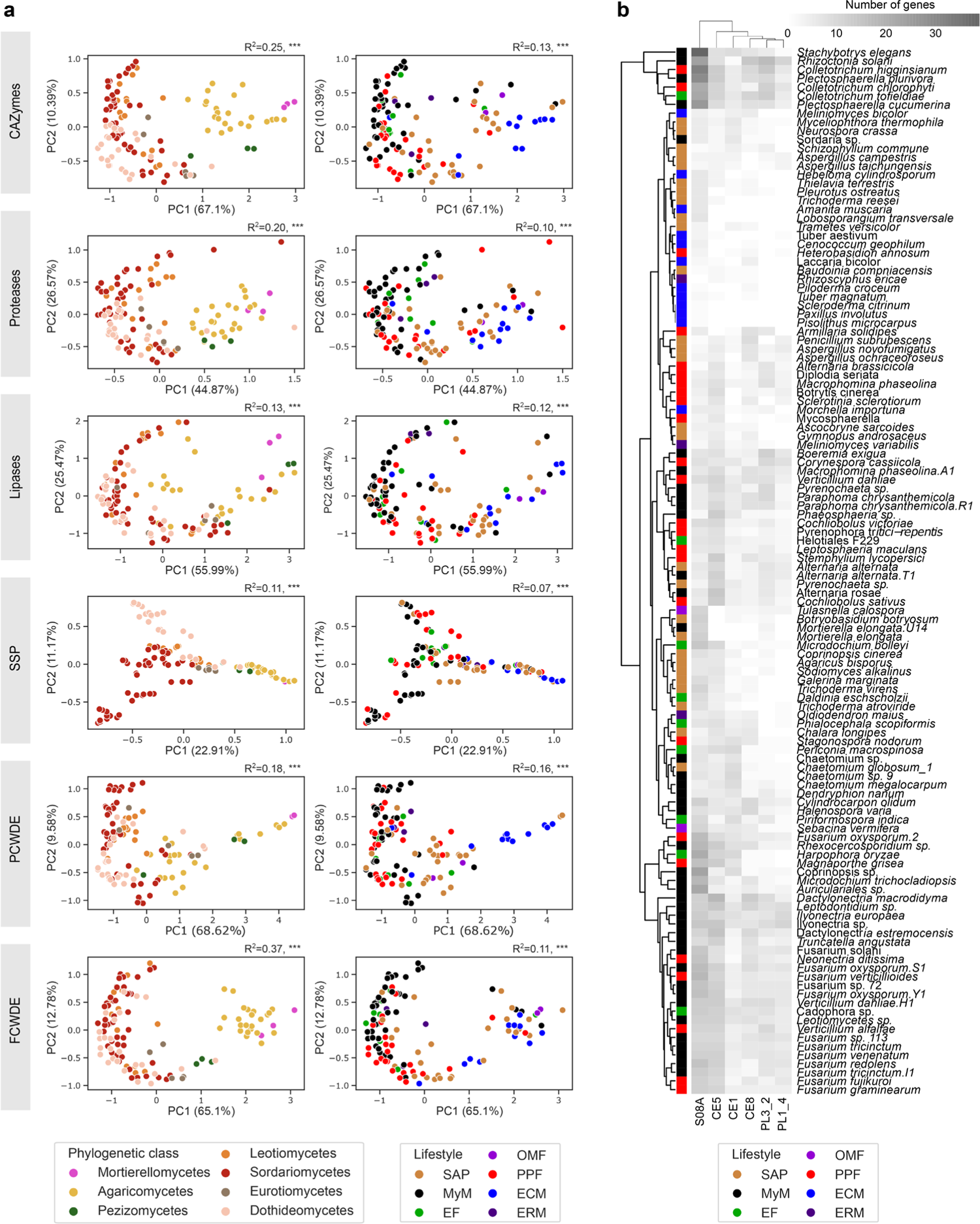
Differential composition in CAZyme, protease, lipase and SSP repertoires according to fungal phylogeny and lifestyle. **a**, Principal component analysis of Jaccard distances calculated on the genomic compositions in subfamilies of CAZymes, proteases, lipases, small secreted proteins (SSP), plant cell wall degrading enzymes (PCWDE) and fungal cell-wall degrading enzymes (FCWDE). Left and right columns of graphs are identical except for the colour coding showing, respectively, fungal phylogeny and fungal lifestyle. Both of these factors significantly explain genomic compositions (PERMANOVA, JaccardMatrix∼Phylogeny+Lifestyle, *P* < 0.05). **b,** Distributions of high loading genes for secreted proteins. S08A: a subfamily S8A secreted serine proteases from proteinase K subfamily. CE: Carbohydrate esterases. PL: Polysaccharide lyases. Colours indicate the ecological style. SAP: Saprotrophs, EF: Endophytic Fungi, PPF: Plant Pathogenic Fungi, ECM: Ectomycorrhiza, ERM: Ericoid Mycorrhiza, OMF: Orchid Mycorrhizal Fungi, MyM: *A. thaliana* mycobiota members. Double clustering heatmap grouping the fungi based on the gene count. Principal components were calculated with theoretically secreted and the total present in the genomes for predicted secretome. High loading genes were determined based on the first three principal components.

**Extended Data Fig. 5:**
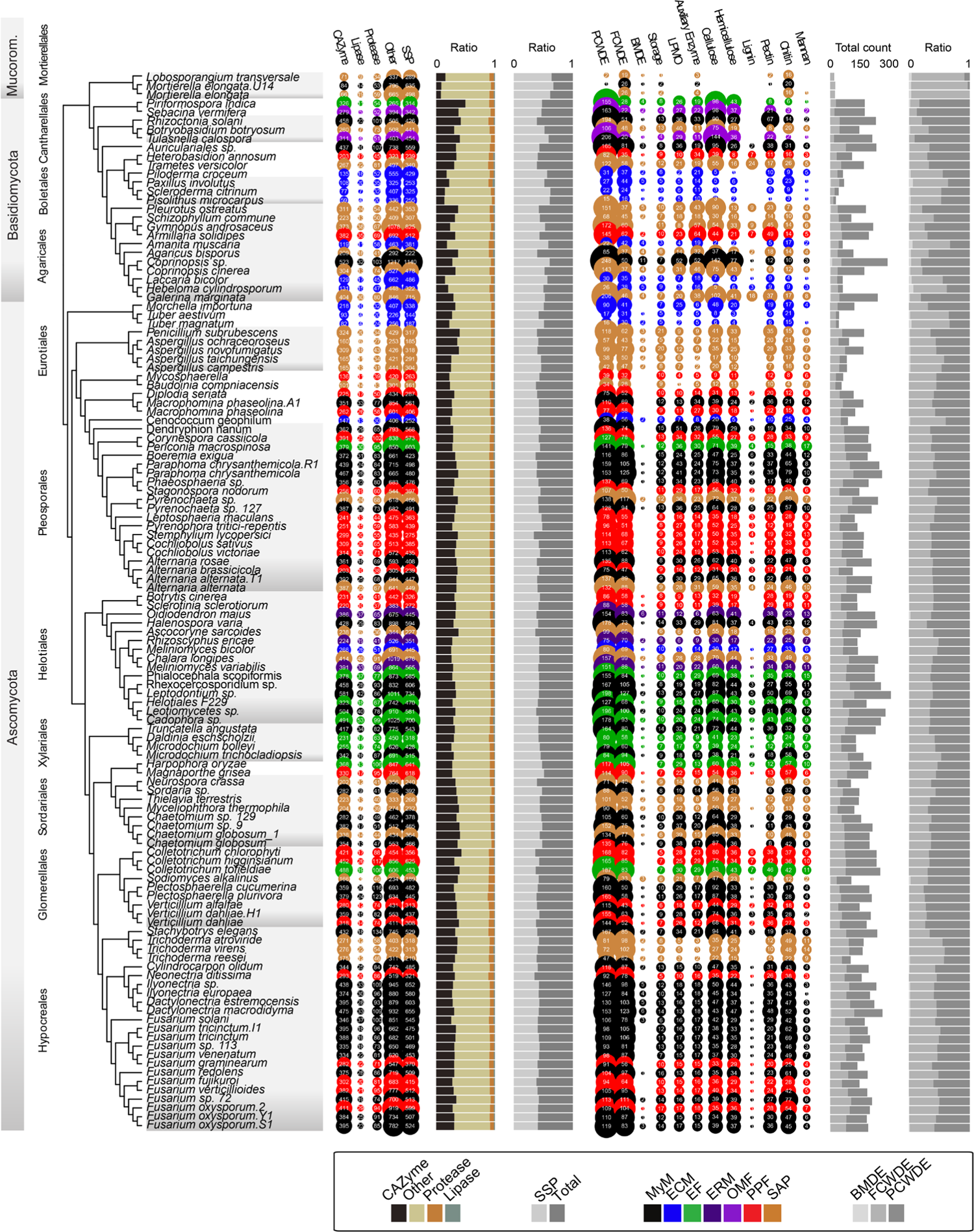
Descriptions and compositions of theoretical fungal secretomes. The first bubble plot (on the left) shows the number of secreted genes for CAZymes, lipases, proteases, and others (i.e., all secreted proteins not in these first three groups). The group SSPs is a subcategory showing the number of small secreted proteins (< 300 aa). The size of bubbles corresponds to the number of genes. The fungi are coloured according to their ecology. The first bar plots (in the middle) represent the ratio of CAZymes, lipases and proteases, to all secreted proteins (left); and the ratio of SSPs among the entire secretome (right). The second bubble plot (on the right) shows CAZyme grouped according to their functions including plant cell-wall degrading enzymes (PCWDE) and fungal cell wall degrading enzymes (FCWDE), peptidoglycans (i.e., bacterial membrane) degrading enzymes (BMDE), trehalose, starch, glycogen degrading enzymes (Storage), lytic polysaccharide monooxygenase (LPMO), substrate-specific enzymes for cellulose, hemicellulose, lignin, and pectin (plant cell walls); chitin, glucan, mannan (fungal cell walls). The second bar plots (far right) show the total count of genes including PCWDE, MCWDE, and BMDE (left); and the proportion of PCWDE, MCWDE, and BMDE (right). SAP: Saprotrophs, EF: Endophytic Fungi, PPF: Plant Pathogenic Fungi, ECM: Ectomycorrhiza, ERM: Ericoid Mycorrhiza, OMF: Orchid Mycorrhizal Fungi, MyM: *A. thaliana* mycobiota members.

**Extended Data Fig. 6:**
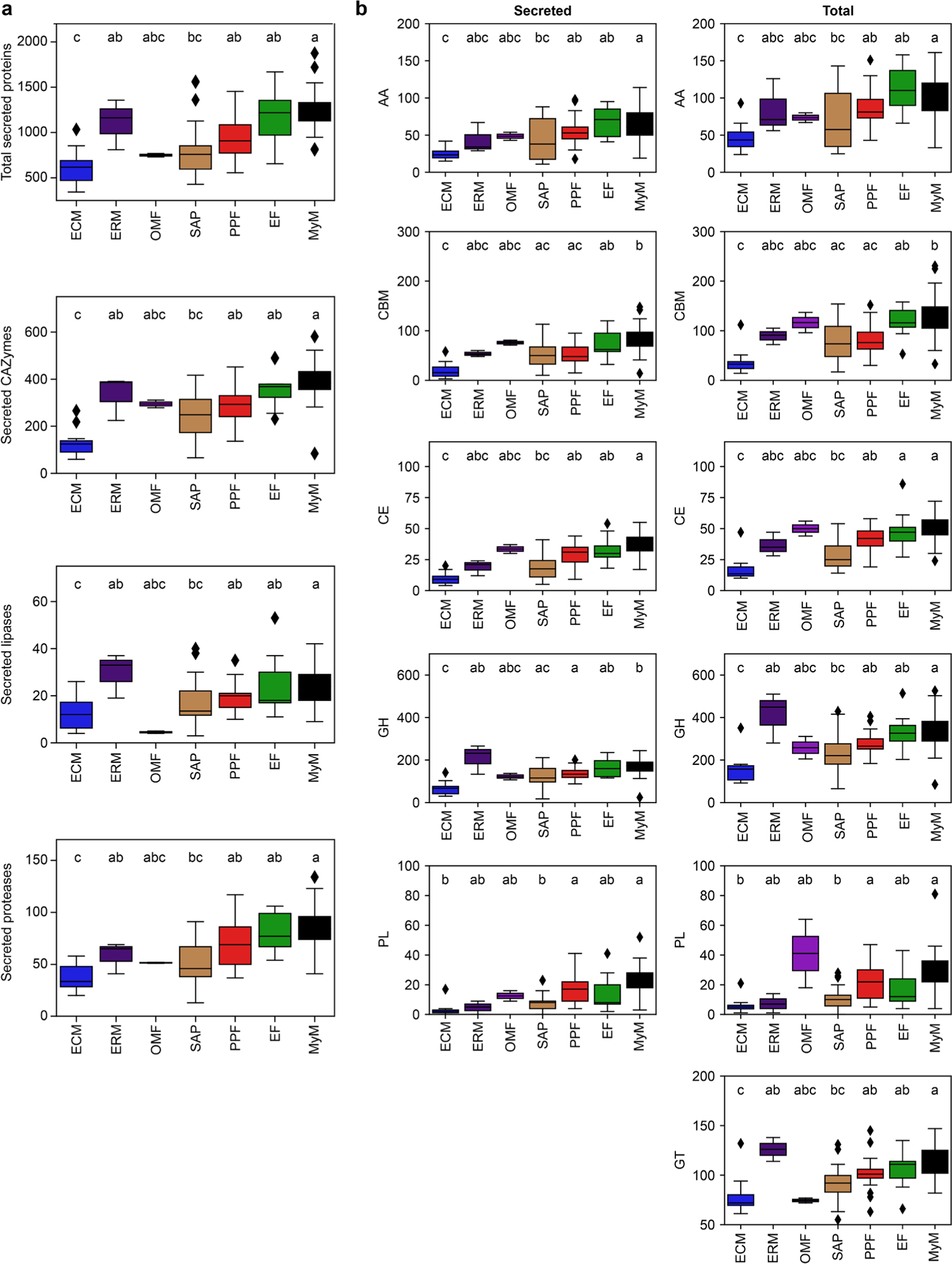
Genomic counts of secreted CAZymes (and subfamilies), proteases and lipases across fungal lifestyles. **a**, Genomic counts of total secreted proteins, and secreted CAZymes, lipases and proteases. ANOVA-statistical testing (Counts∼PhylogenyPCs+Lifestyle, **Methods**) identified both phylogeny and lifestyles as having an effect on genomic contents; letters result from post-hoc TukeyHSD testing. **b**, Gene counts of CAZyme families (AA: Auxiliary Activities, CBM: Carbohydrate-Binding Modules, CE: Carbohydrate Esterases, GH: Glycoside Hydrolases, PL: Polysaccharide Lyases), predicted as secreted (extracellular, left) and total (intra and extracellular, right). Statistical testing with Kruskal-Wallis (Counts∼Lifestyle) identified lifestyle as having an effect on genome contents. Letters result from post-hoc testing with a Dunn test. SAP: Saprotrophs, EF: Endophytic Fungi, PPF: Plant Pathogenic Fungi, ECM: Ectomycorrhiza, ERM: Ericoid Mycorrhiza, OMF: Orchid Mycorrhizal Fungi, MyM: *A. thaliana* mycobiota members.

**Extended Data Fig. 7:**
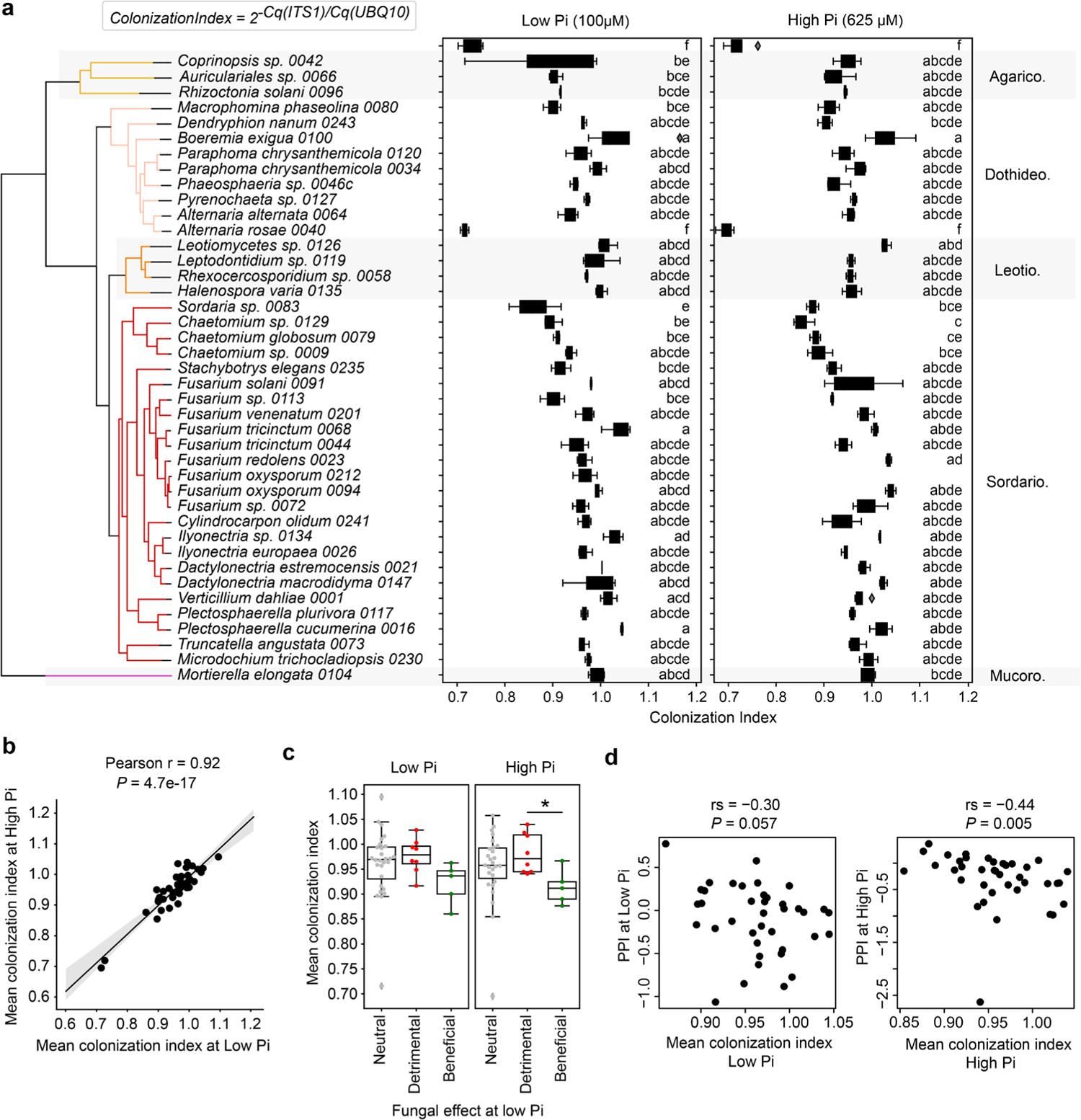
Fungal colonization of roots after 28 days of culture in mono-association. **a)** Fungal colonization of plant roots mono-inoculated with different mycobiota members, estimated by quantitative PCR (**Methods**). Statistical difference across treatments was identified by ANOVA (ColonizationIndex∼Treatment), and post-hoc testing was performed with TukeyHSD. **b**, Pearson correlation between colonization indexes at low and high Pi concentrations (*P* < 0.05). **c**, Differences in colonization indices between neutral, beneficial, and detrimental fungi at low Pi. Significant differences across fungal groups were identified by ANOVA and TukeyHSD. **d**, Correlation between plant performance index and mean colonization index at low Pi (left) and high Pi (right) (n = 41, Spearmańs rank correlation).

**Extended Data Fig. 8:**
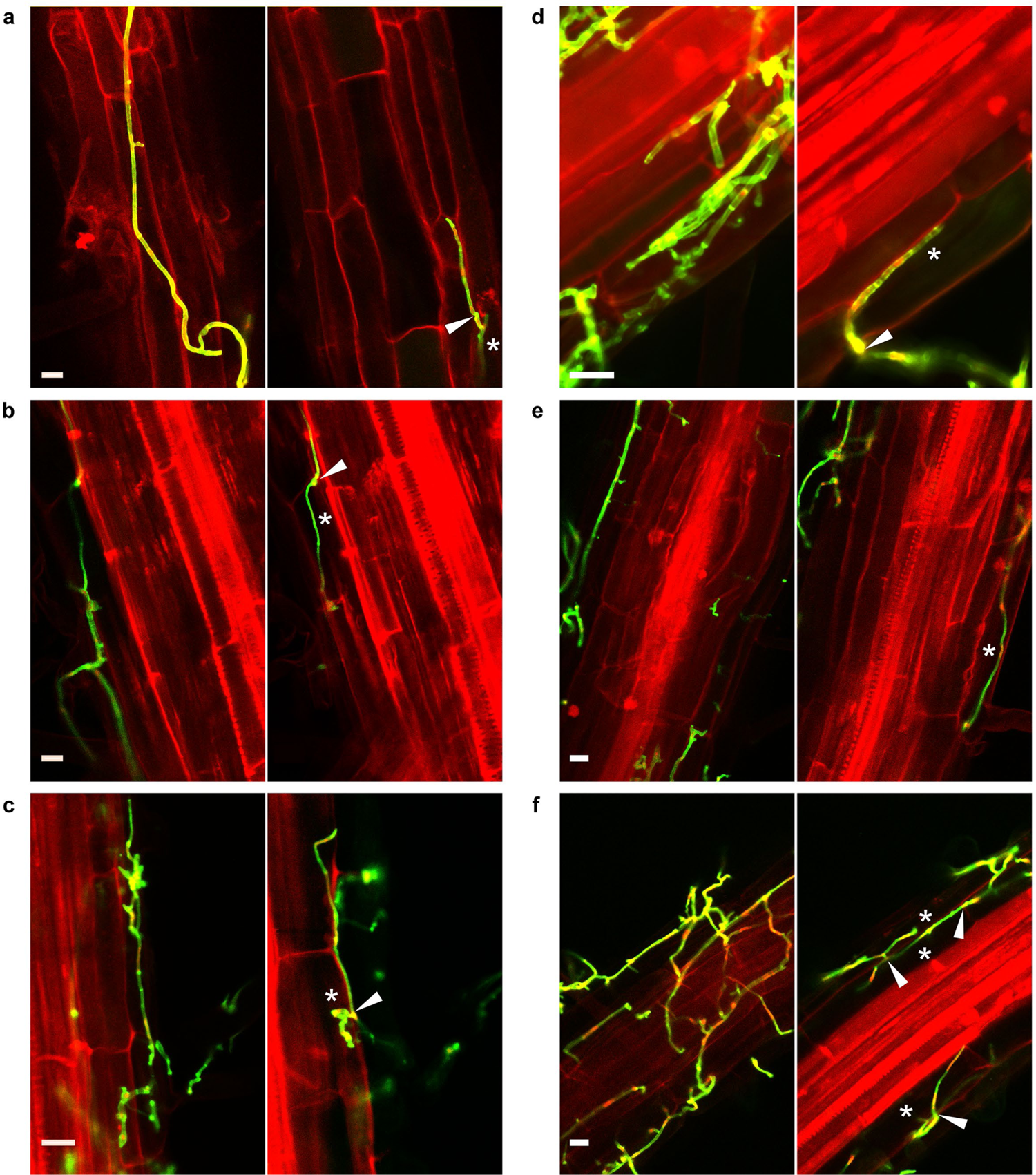
Confocal imaging of root surface and epidermis colonization by mycobiota members. Roots grown for 4 weeks in mono-association with six diverse fungi, double-stained with Propidium Iodide and Wheat Germ Agglutinin coupled to fluorophore CF488 (WGA-488; Biotium), imaged by confocal microscopy. Left and right picture belong to a single z-stack, respectively focusing on the root surface and the root endosphere where colonization of epidermal cells can be observed. Arrows indicate penetration sites and asterisks infected root cells. **a,** *Cs* = *Chaetomium sp. 0009.* **b**, *Mp* = *Macrophomina phaseolina 0080. **c**, Pc= Paraphoma chrysanthemicola 0120. **d***, *Ps* = *Phaeosphaeria sp. 0046c.* **e**,*Ta* = *Truncatella angustata 0073* **f**, *Hv* = *Halenospora varia 0135*.

**Extended Data Fig. 9:**
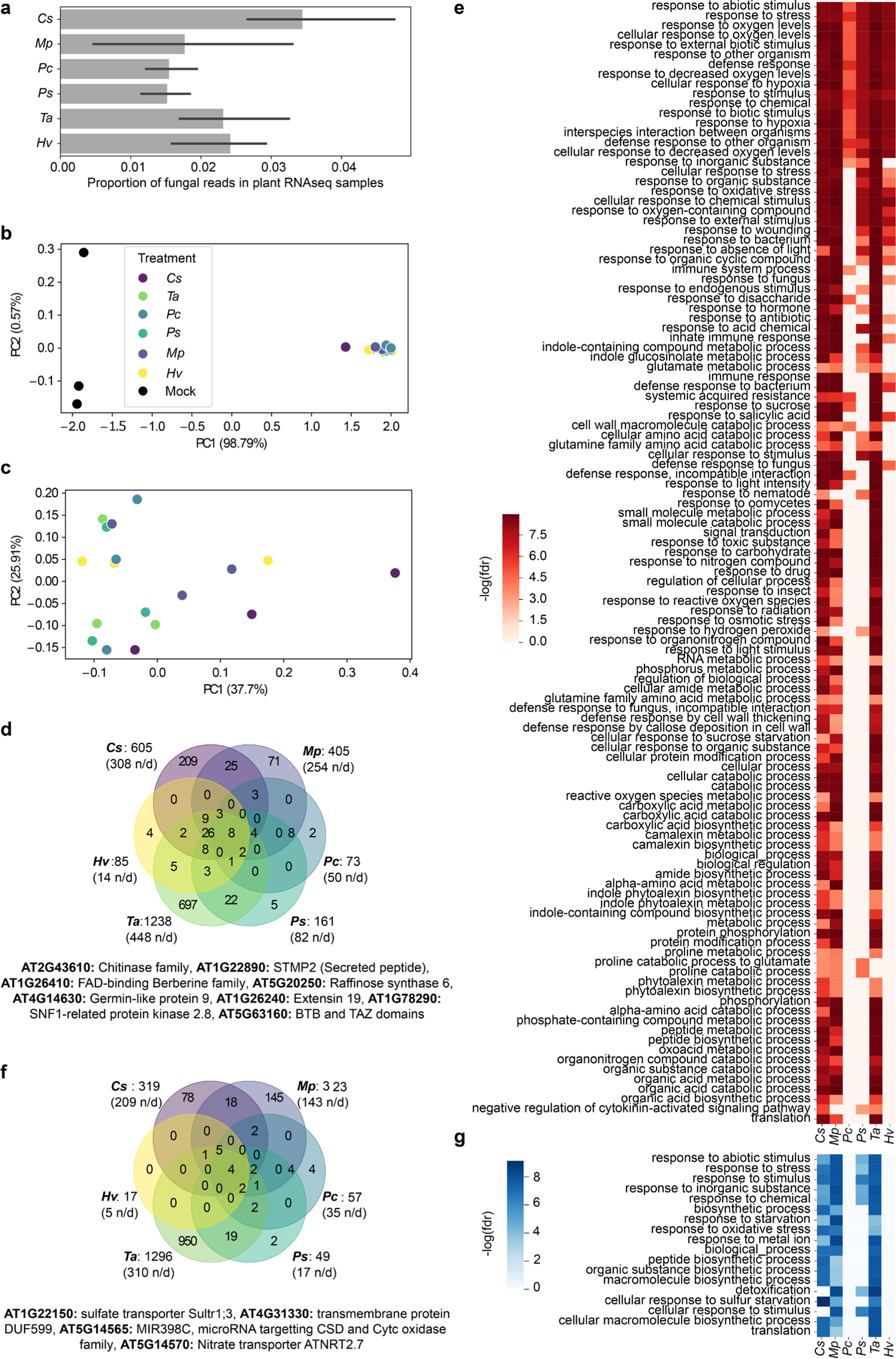
*Arabidopsis thaliana* transcriptional reprogramming upon colonization by different root mycobiota members. **a**, Proportion of reads in RNA-Seq samples mapped on fungal genomes. **b**, Principal Component Analysis of Bray-Curtis distances calculated over *A. thaliana* gene read counts. **c**, Principal Component Analysis of Bray-Curtis distances calculated over *A. thaliana* gene read counts, excluding mock-treated samples to reveal sample differences due to the different fungi. *Cs* = *Chaetomium sp. 0009*, *Mp* = *Macrophomina phaseolina 0080*, *Pc* = *Paraphoma chrysantemicola 0034*, *Ps* = *Phaeosphaeria sp. 0046c*, *Ta* = *Truncatella angustata 0073*, *Hv* = *Halenospora varia 0135.* **d**, Venn diagram showing *A. thaliana* commonly over-expressed genes in response to fungal inoculations. On the right is the list of genes over-expressed in response to all six fungi. **e**, Independent GO enrichment analyses performed on the *Arabidopsis* genes over-expressed in response to each fungus (GOATOOLS, FDR < 0.05). **f**, Venn diagram showing *A. thaliana* commonly under-expressed genes in response to fungal inoculations. On the right is the list of genes under-expressed in response to all six fungi. **g**, Independent GO enrichment analyses performed on the *Arabidopsis* genes under-expressed in response to each fungus (GOATOOLS, FDR < 0.05).

**Extended Data Fig. 10:**
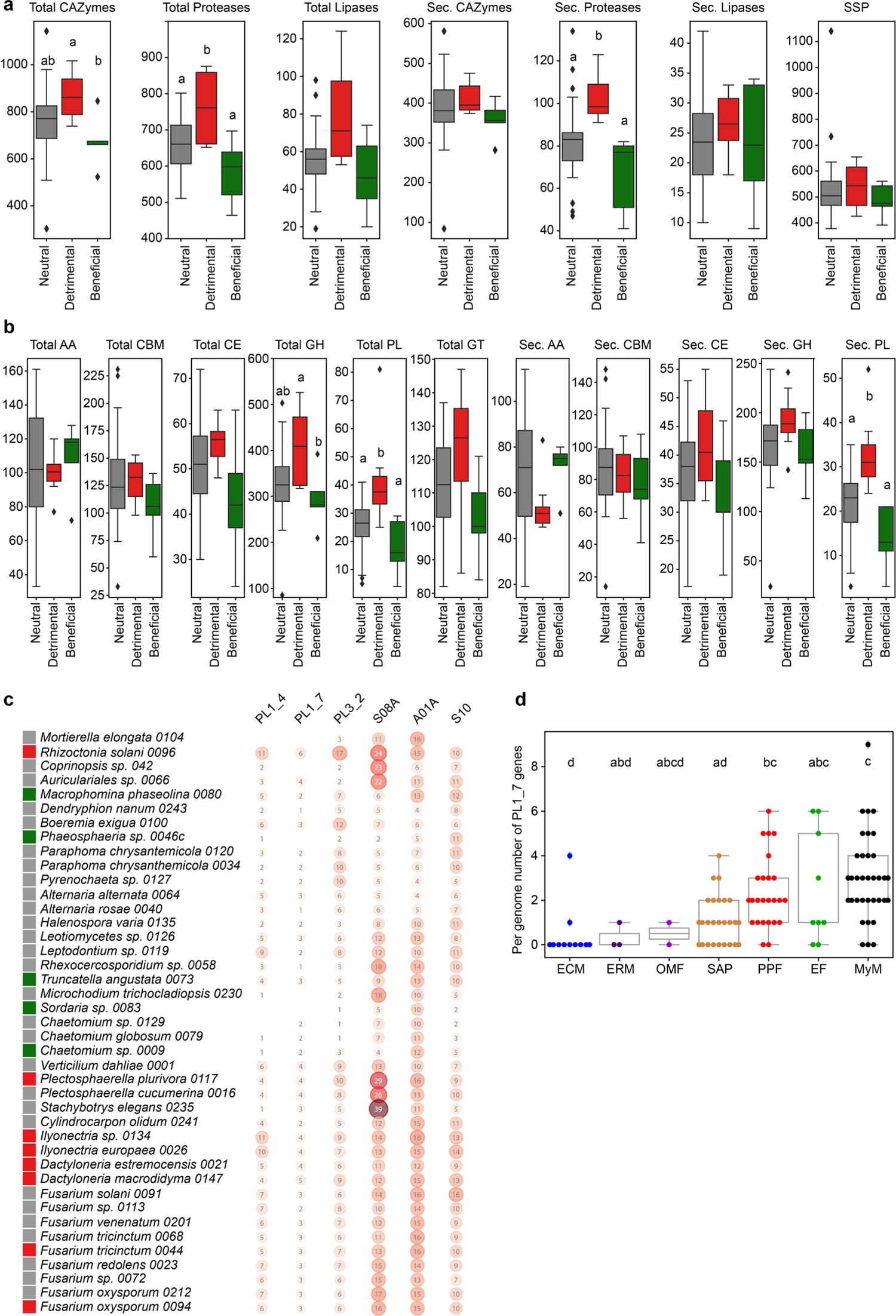
Genomic signatures in polysaccharide lyase repertoires explain lifestyle differentiation among root mycobiota members. **a**, Distribution of genes encoding secreted or total (secreted with intracellular) proteins in 41 endophytic fungi. a) CAZymes, lipases, protease, SSPs. **b,** CAZyme family. Different letters indicate significant difference (FDR adjusted *P* < 0.05; Kruskal Wallis test). Beneficial (n = 5), neutral (26), pathogenic (10). **c,** Key secreted protein coding genes discriminating fungal lifestyles of 41 endophytic fungi. Three fungal lifestyles are depicted in colour. The selected genes coding for secreted polysaccharide lyases (PLs) and proteases discriminate between pathogenic, neutral, and beneficial endophytic fungi. Fungal taxa are displayed according to the phylogenetic order. Bubbles with numbers contain the number of genes. **d,** Comparative genomics of PL1_7, showing the number of PL1_7 genes in the genomes associated to different lifestyles. Statistics were performed using an ANOVA test and a TukeyHSD post-hoc test.

## Supplementary Tables

**Supplementary Table 1:** Description of the 41 newly-sequenced strains of *A. thaliana* root mycobiota members

**Supplementary Table 2:** Description of the comparative genomics dataset, comprising our 41 newly-sequenced strains and 79 published fungal genomes

**Supplementary Table 3:** Orthogroups that segregate best endophytes and mycobiota members from other fungi. **a**, Description of the 84 orthogroups. **b**, GO enrichment analysis performed on the 84 orthogroups. **c**, Coexpression of the 84 gene families in fungi, according to STRING^29^.

**Supplementary Table 4:** Differential fungal gene expression in planta *vs.* on medium.

**Supplementary Table 5:** Differential host gene expression inoculated with fungi *versus* mock-treated.

**Supplementary Table 6:** Description of the 11 orthogroups segregating detrimental mycobiota members from others.

